# Triple-Omics Analysis Unveiled the Neuroprotective Mechanisms of *Akkermansia muciniphila* in an Environmental Exposure-Linked Mouse Model of Alzheimer’s Disease

**DOI:** 10.64898/2026.09.30.755812

**Authors:** Zachary Kowash, Joe Lim, Andrew J. Annalora, Hao Wang, Julia Yue Cui

## Abstract

Cadmium (Cd) is an emerging environmental risk factor for cognitive decline and Alzheimer’s disease, and few interventions limit its effects on the brain. *Akkermansia muciniphila*, a human gut commensal, is neuroprotective through mechanisms that remain unclear. Building on our finding that *A. muciniphila* prevented Cd-induced memory deficits without reducing brain Cd burden (Wang et al., 2026a), we tested whether it reprograms Cd-disrupted hippocampal RNA regulation. Male C57BL/6J mice received *A. muciniphila* (1×10⁹ CFU, oral gavage, 5 days/week) before and throughout 9 weeks of 3 mg/L CdCl₂ exposure; hippocampi were analyzed by RNA-seq, m⁶A epitranscriptomic profiling, and rMATS-based alternative-splicing analysis. Cd disrupted hippocampal neuropeptide signaling, neuronal excitability, and synaptic plasticity. Co-exposure with *A. muciniphila* reversed signatures enriched for Nrf2-centered cytoprotective signaling and opposed m⁶A and splicing changes across more than a thousand genes, largely without changes in total mRNA abundance. Restored events converged on machinery governing local translation, calcium-dependent plasticity, and synaptic transmission. Projected onto predicted protein structures, most introduced a premature termination codon predicted to trigger nonsense-mediated decay or edited short loop and linker segments rather than rebuilding folded domains, regulation largely invisible to gene-level expression analysis. Among these, paired events amplifying semaphorin-plexin signaling, an axis linked to oligodendrocyte injury and memory consolidation, were reversed. These findings identify restoration of hippocampal transcript fate and splice-isoform selection as a mechanism potentially underlying *A. muciniphila*-mediated protection against Cd neurotoxicity, and establish post-transcriptional RNA regulation as a gut-brain target for mitigating environmentally driven neurotoxicity.

**IMPACT STATEMENT:** Cadmium is a widespread environmental contaminant associated with cognitive decline and Alzheimer’s disease, and few interventions are available to limit its effects on the brain. This study shows that supplementation with the gut commensal *Akkermansia muciniphila* reverses Cd-induced changes in hippocampal m⁶A methylation and splice-isoform selection, largely without altering total transcript abundance. These results identify post-transcriptional RNA regulation as a gut-brain target for mitigating environmentally driven neurotoxicity, and show that conventional gene-expression analysis alone would miss most of the protective response.

## INTRODUCTION

Epitranscriptomic regulation, particularly N⁶-methyladenosine (m⁶A), supports neuronal development, plasticity, and repair, and its disruption contributes to neurological disease. m⁶A dynamically regulates neuronal gene expression, dendritic structure, synapse formation, and memory, and its patterns vary across neurodevelopment, brain region, and aging; dysregulated m⁶A signaling and reduced m⁶A modification of Alzheimer’s disease (AD)-related transcripts are associated with AD pathology and neuronal loss (Shafik et al., 2021; Hashmi et al., 2024). The machinery is functionally decisive: methyltransferase-like 3 (METTL3) promotes hippocampal long-term memory consolidation (Zhang et al., 2018), the reader YTH N⁶-methyladenosine RNA-binding protein 1 (YTHDF1) facilitates hippocampus-dependent learning (Shi et al., 2018), and the eraser fat mass and obesity-associated protein (FTO) regulates m⁶A-dependent memory formation (Walters et al., 2017). Alternative splicing, which is itself regulated by m⁶A (Xiao et al., 2016; Wang CX et al., 2018), is likewise essential for neuronal identity, synaptic signaling, and stress adaptation. In sporadic AD, affected regions show altered splicing of tau exon 10 and related transcripts (Glatz et al., 2006), and human temporal cortex carries AD-specific splicing changes distinct from normal aging (Tollervey et al., 2011). Reducing T-cell intracellular antigen 1 (TIA1) partially restores normal splicing and promotes neuroprotection in tauopathy mice (Apicco et al., 2019), while loss of nuclear TAR DNA-binding protein 43 (TDP-43) de-represses cryptic exons in stathmin-2 (STMN2) and unc-13 homolog A (UNC13A) (Agra Almeida Quadros et al., 2024; Ma et al., 2022). Together, m⁶A and splicing form a coupled regulatory layer whose disruption promotes synaptic dysfunction and neurodegeneration.

The gut-brain axis is increasingly recognized as a regulator of neurological function through metabolic, immune, intestinal-barrier, and neurotransmitter pathways (Dempsey et al., 2019). *Akkermansia muciniphila* is a mucin-degrading commensal that constitutes approximately 1–3% of the gut microbiota in healthy adults (Collado et al., 2007; Derrien et al., 2008). Its neurological benefits span multiple models: reduced cerebral amyloid-beta and improved memory in an AD mouse model (Ou et al., 2020), improved cognition in aged mice through suppression of interleukin-6-associated inflammation (Zhu et al., 2023), preserved dopaminergic neurons in a Parkinson’s disease model (Qiao et al., 2024), and mediation of the antiseizure effect of the ketogenic diet through hippocampal γ-aminobutyric acid and glutamate balance (Olson et al., 2018). These findings support a therapeutic role for *A. muciniphila* in neurological health through the gut-brain axis.

Cadmium (Cd) is an emerging environmental risk factor for AD, with exposure concerns spanning early brain development through later-life neurodegeneration, and epidemiological evidence increasingly implicates environmental metal exposures in AD risk (Armstrong, 2019). In older U.S. adults, higher blood cadmium was associated with greater AD mortality (Min and Min, 2016), a finding reproduced using urinary Cd as a marker of cumulative exposure (Peng et al., 2017), and higher blood Cd has been linked to poorer cognition and greater cognitive decline (Liu et al., 2021). Preclinical studies provide biological plausibility, showing that Cd impairs hippocampal neurogenesis and promotes oxidative stress, neuroinflammation, amyloid-beta accumulation, tau pathology, and cognitive deficits in genetically susceptible AD models (Ali et al., 2021; Liu et al., 2023; Li et al., 2024a). Cd also disrupts RNA regulation directly: maternal Cd exposure causes hippocampal m⁶A hypomethylation and suppresses YTHDF2-mediated degradation of *Bax* mRNA, increasing neuronal apoptosis and cognitive deficits in offspring (Wang et al., 2026b), and Cd disrupts RNA-splicing pathways relevant to neuronal injury (Del Pino et al., 2016; Wu et al., 2019). We recently demonstrated that oral supplementation with human fecal microbiome-derived *A. muciniphila* prevented cognitive impairment in Cd-exposed AD mice across 9 weeks of exposure at a human body burden-relevant dose, without reducing brain Cd concentrations (Wang et al., 2026a) — indicating protection through mechanisms independent of Cd disposition in the brain.

Although Cd neurotoxicity and the neuroprotective effects of *A. muciniphila* are each established, the mechanism underlying this protection remains unknown. It is unknown whether *A. muciniphila* protects against Cd-induced cognitive impairment by reprogramming the hippocampal epitranscriptomic and alternative-splicing landscapes to restore disease-relevant gene-expression networks. Addressing this gap would define a previously unexplored microbiome-RNA regulatory mechanism linking an environmental toxicant to neurodegenerative disease. We therefore tested the hypothesis that *A. muciniphila* reprograms the hippocampal m⁶A and alternative-splicing landscapes to restore neuroprotective gene-expression networks, using a triple-omics approach integrating m⁶A epitranscriptomic profiling, rMATS-based alternative-splicing analysis, and RNA-seq transcriptome-wide expression analysis.

## MATERIALS AND METHODS

### Chemicals, reagents, and bacteria

Cadmium chloride (CdCl₂; catalog no. 202980) was obtained from MilliporeSigma (Burlington, MA), and acetic acid was sourced from Thermo Fisher Scientific (Fair Lawn, NJ). The following analytical standards and derivatization reagents were purchased from Sigma-Aldrich (St. Louis, MO): propionic acid, isobutyric acid, butyric acid, 2-methylbutyric acid, isovaleric acid, valeric acid, 2-methylpentanoic acid, 3-methylpentanoic acid, isocaproic acid, caproic acid, 2-methylhexanoic acid, 4-methylhexanoic acid, heptanoic acid, hexanoic acid-6,6,6-d₃ internal standard, N-tert-butyldimethylsilyl-N-methyltrifluoroacetamide (MTBSTFA), and methoxyamine hydrochloride. Unless otherwise specified, all remaining chemicals were also obtained from Sigma-Aldrich. *A. muciniphila* (BAA-835) was obtained from ATCC, Manassas, VA.

### Animals and experimental exposures

Six-week-old male C57BL/6J mice were obtained from The Jackson Laboratory (Bar Harbor, ME). Animals were housed five per cage under standard laboratory conditions with a 12-hour light/12-hour dark cycle and provided unrestricted access to food and water. Mice were fed PicoLab Rodent Diet 20 (LabDiet, St. Louis, MO). Drinking water consisted of reverse-osmosis–purified tap water acidified with 2.4–2.8% HCl and sterilized by autoclaving.

Beginning at eight weeks of age, mice were administered *A. muciniphila* or the corresponding vehicle by oral gavage. Treatments were delivered at 10 mL/kg once daily for five consecutive days each week, Monday through Friday. The bacterial suspension contained 1 × 10⁹ colony-forming units in 0.5 g/L L-cysteine, which was included to help preserve bacterial viability under anaerobic conditions. Administration of *A. muciniphila* began one week before Cd exposure. Following the initial week of *A. muciniphila* treatment, mice received either standard drinking water or drinking water containing 3 mg/L CdCl₂ –a concentration selected to produce blood Cd concentrations close to those reported in U.S. smokers (Wang H et al., 2018) – for nine weeks beginning at nine weeks of age (Fig. 1A). Treatment with *A. muciniphila* or vehicle was maintained throughout the Cd exposure period, resulting in a total treatment duration of 10 weeks. Thus, in total there were four experimental groups: vehicle control (Ctrl), *A. muciniphila alone* (Akk), cadmium alone (Cd), and combined A. muciniphila and cadmium exposure (Akk+Cd). At the conclusion of the experiment, week 10, mice were euthanized and hippocampus was collected for subsequent analyses. Animal husbandry and experimental procedures were approved by the Institutional Animal Care and Use Committees of the University of Washington and the University of Michigan.

**Figure 1.**
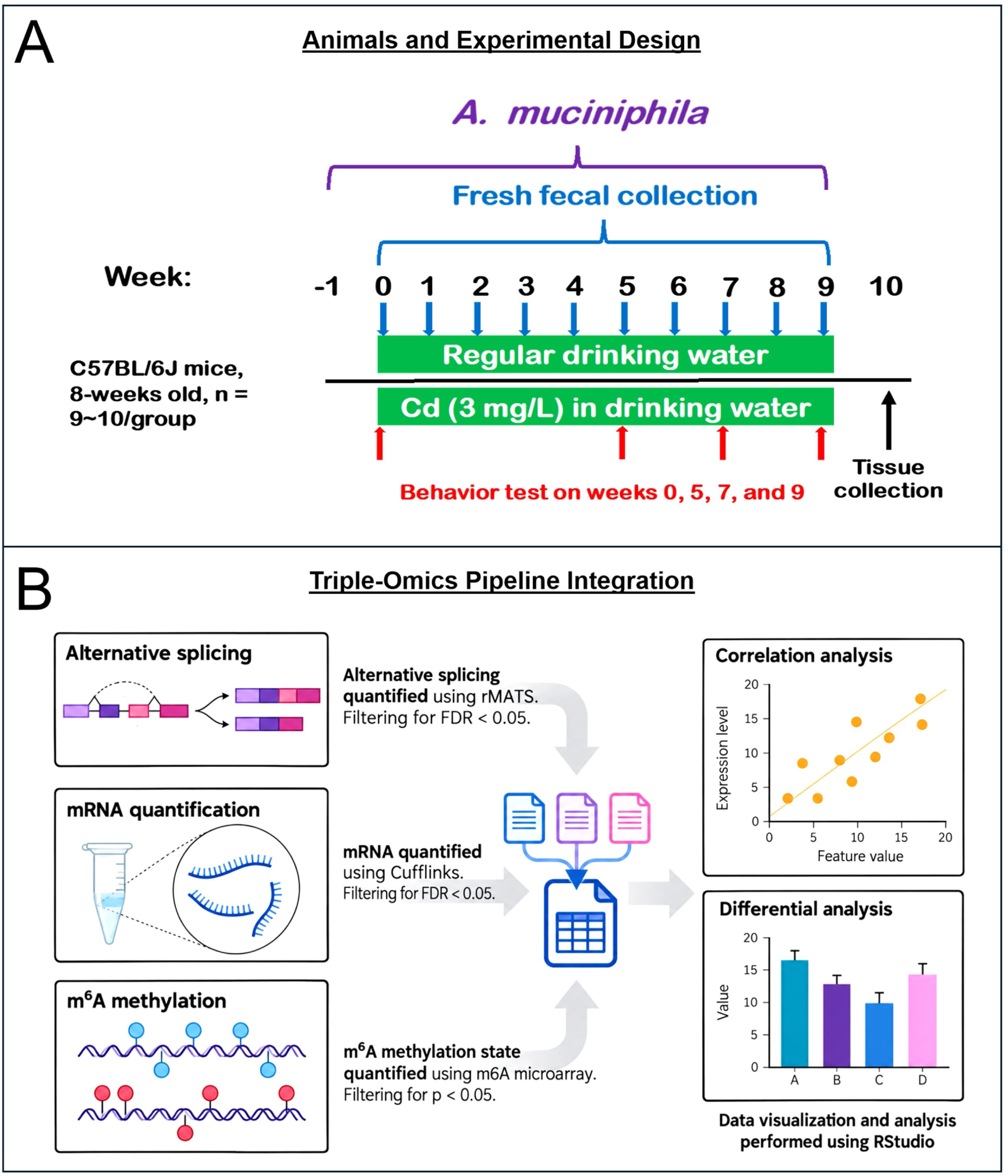
Experimental design and hippocampal triple-omics analysis workflow. A. Experimental design for cadmium (Cd) exposure and *Akkermansia muciniphila* (Akk) supplementation in male C57BL/6J mice. Akk supplementation was initiated one week before Cd exposure and continued throughout the study. CdCl2 was administered in drinking water at 3 mg/L, with longitudinal fecal collection and tissue collection at the end of the exposure period (Wang et al., 2026a). **B.** Schematic of the hippocampal triple-omics workflow integrating RNA-seq-based mRNA expression, m⁶A epitranscriptomic profiling, and rMATS-based alternative-splicing (AS) analysis, followed by integrated statistical analysis and visualization in R.

### Total RNA Isolation

Total RNA was extracted from hippocampal samples using RNA-Bee reagent (Tel-Test Inc., Friendswood, TX) according to the manufacturer’s protocol RNA concentrations were quantified using NanoDrop 1000 Spectrophotometer (Thermo Scientific, Waltham, MA) at 260 nm. The integrity of total RNA samples was evaluated by formaldehyde-agarose gel electrophoresis with visualization of 18S and 28S rRNA bands under UV light, and by an Agilent 2100 Bioanalyzer (Agilent Technologies Inc., Santa Clara, CA). Samples with RNA integrity numbers above 8.0 were used for RNA-Seq. Samples were then subjected to bulk RNA sequencing.

### Bulk RNA-sequencing analysis

Bulk RNA-sequencing was performed by Novogene (2×150 bp paired ends, 20 million mapped reads per sample), and fastq files were aligned to the mouse reference genome (GRCm38/mm10) using HISAT2. Transcript abundance was calculated using Cuffllinks and differential expression was assessed using Cuffdiff. Gene-level log₂ fold-change values were obtained from a precomputed expression matrix and matched to Cuffdiff q-values using gene symbols. Group-level mean FPKM values and standard errors were calculated from the same expression matrix for visualization. Pathway analyses were conducted using Enrichr (https://maayanlab.cloud/Enrichr/) (FDR-BH < 0.05).

### m⁶A RNA-methylation analysis

N⁶-methyladenosine (m⁶A) RNA profiling was performed using the Arraystar Mouse Epitranscriptomic Microarray (Arraystar Inc., Rockville, MD). Total RNA was subjected to immunoprecipitation with an anti-m⁶A antibody to separate m⁶A-enriched RNA from the unbound RNA fraction. The immunoprecipitated and supernatant RNA fractions were converted to cRNA, labeled with Cy5 and Cy3, respectively, and co-hybridized to the microarray. Normalized fluorescence intensities from the two channels were used to estimate transcript-level m⁶A enrichment and identify differentially methylated transcripts (*p* < 0.05). Positive m⁶A log₂ fold-change values were classified as hypermethylation, whereas negative values were classified as hypomethylation. Fold-change values were log₂-transformed after adding a pseudocount of 10⁻⁶ to prevent undefined values for zero measurements. When multiple transcripts mapped to the same gene, the transcript with the smallest p-value was retained to generate a single gene-level record.

### Alternative-splicing analysis

Alternative-splicing events were quantified using replicate Multivariate Analysis of Transcript Splicing (rMATS). Five major event types were evaluated: skipped exons (SE), alternative 5′ splice sites (A5SS), alternative 3′ splice sites (A3SS), mutually exclusive exons (MXE), and retained introns (RI). Events with a *p* < 0.05 were considered significant. To define a high-confidence subset of restoration events, splicing decisions were additionally required to meet a Benjamini–Hochberg FDR < 0.05 in both the Cd vs. Vehicle and Cd+Akk vs. Cd comparisons, to change in opposite directions between those comparisons, to show a Cd effect of at least 10% PSI, and to show no residual departure from the vehicle baseline (Cd+Akk vs. Vehicle, FDR ≥ 0.05). The study design comprises four arms (vehicle, A. muciniphila alone, Cd, and Cd+A. muciniphila), and rMATS was additionally run for the Cd+Akk vs. Vehicle comparison, so the position of the co-treated group relative to vehicle is taken from that measured contrast and carries its own FDR. This fourth contrast was run for the alternative-splicing layer only. Because ΔPSI is a difference of group means, this value is identical to the sum ΔPSI(Cd vs. Vehicle) + ΔPSI(Cd+Akk vs. Cd) wherever no replicate is dropped for insufficient coverage, which holds for every event in the high-confidence set.

For genes containing multiple significant splicing events, the gene-level splicing effect was summarized as the mean inclusion-level difference or change in percent-spliced-in value (ΔPSI), across all significant events. The minimum event-level p-value was retained as the corresponding gene-level significance measure. Genes with a positive mean ΔPSI were classified as exhibiting increased exon inclusion, whereas genes with a negative mean ΔPSI were classified as exhibiting decreased exon inclusion. For transcriptome-wide correlation analyses, all genes evaluated by rMATS were included regardless of statistical significance.

### Structure-function analysis of restored splicing events

To interpret the protein-level consequence of the high-confidence restored events, each rMATS event was mapped to the corresponding RefSeq exon and the affected nucleotide segment was resolved on the annotated transcript. Reading frame was determined from the length of that segment. Segments whose length was not a multiple of three were classified as frameshifting and predicted to introduce a premature termination codon, and were therefore treated as acting through nonsense-mediated decay rather than altering protein sequence locally. In-frame segments were translated and mapped to residue ranges on the corresponding UniProt canonical sequence (UniProt Consortium, 2025).

Each in-frame residue range was then characterized on the AlphaFold model of that protein, retrieved from the AlphaFold Protein Structure Database (Jumper et al., 2021; Varadi et al., 2024). Three model-derived features were used, together with the position of the segment within the chain: secondary-structure assignment across the segment; relative burial, calculated as the mean number of Cα atoms within 10 Å of each segment Cα and normalized to the median value for that protein, with a normalized value ≥ 1.15 taken as buried; and the per-residue model confidence score (pLDDT), with values below 50 treated as disordered and values of 70 or above as ordered. On this basis each segment was assigned a structural character: surface loop, disordered insert, buried helix or strand, or terminal segment.

Two annotation layers were then intersected with the same residue range. Curated UniProt sequence features (active and binding sites, cofactor and metal sites, post-translational modification sites, disulfide bonds, signal, transit and transmembrane segments, DNA-binding regions, and annotated interaction regions) were used to identify segments that remove a specific functional element (UniProt Consortium, 2025). InterPro domain assignments (Blum et al., 2025) were used to describe the position of the segment relative to domain architecture, distinguishing removal of a complete modular domain, a domain-internal deletion, a domain-boundary or inter-domain linker segment, and a segment lying outside annotated domains. Events were assigned to the four classes shown in Figure 9A in the following order of precedence: predicted nonsense-mediated decay; removal of an entire structural domain; an in-frame edit overlapping an annotated functional element; and an in-frame edit confined to a loop or disordered segment. Each event was additionally rendered in its transcript and reading-frame context using the cassette exon visualization method described previously (CeVip; Annalora et al., 2020), and every structure-function call reported here was inspected in that view. This exon-level assessment follows the approach we previously developed for the nuclear receptor and cytochrome P450 superfamilies, in which the predicted structural consequence of an alternatively spliced exon, rather than its statistical rank, is used to judge which isoforms are worth targeting (Annalora et al., 2020; Lidberg et al., 2021). Structural interpretations reported here are predictions from transcript-level measurements and AlphaFold models, and were not verified experimentally.

### Triple-omics integration and statistical analysis

RNA-seq, m⁶A epitranscriptomic, and alternative-splicing (AS) datasets from mouse hippocampus were integrated across three pairwise comparisons: Cd versus Vehicle, Akkermansia muciniphila (Akk) versus Vehicle, and Cd+Akk versus Cd, as shown in **Fig. 1B**. A fourth comparison, Cd+Akk versus Vehicle, was generated for the alternative-splicing layer only, where it is used solely to test whether a restored event retains a residual departure from the vehicle baseline; the corresponding contrast was not generated for the RNA-seq or m⁶A datasets. The design is therefore not fully balanced across the three layers, and every cross-layer comparison reported here is restricted to the three contrasts common to all of them. Because the study was designed as an exploratory, hypothesis-generating multi-omics analysis, statistical significance was defined using the *p* < 0.05 threshold. This is because multiple-testing correction can be highly stringent in high-dimensional m⁶A and AS datasets, in which thousands of transcript-or event-level features are tested, and may exclude moderate but potentially biologically informative changes that become meaningful when evaluated across multiple molecular layers. Accordingly, p values were applied consistently across datasets to maximize sensitivity for candidate discovery. Gene symbols were standardized to title case across the bulk RNA-seq, m⁶A, and AS datasets before integration. Computationally predicted or incompletely annotated identifiers, including Gm genes, Riken loci, LOC loci, numerical-prefix identifiers, and unannotated Ensembl identifiers, were excluded from enrichment analyses and graphical summaries. Cross-comparison genes were defined as genes that were differentially expressed in both the Cd vs. Control and Cd+Akk vs. Cd comparisons. These genes represented Cd-responsive transcripts that were also significantly modified by *A. muciniphila* co-treatment.

Pairwise relationships among molecular layers were assessed separately for each treatment comparison using Spearman rank correlations. The evaluated pairs were RNA log₂ fold change versus m⁶A log₂ fold change, RNA log₂ fold change versus ΔPSI, and m⁶A log₂ fold change versus ΔPSI. Correlations were calculated using all genes with finite values for both measurements, without restricting the analysis to statistically significant genes. Spearman correlation coefficients and two-sided p-values were calculated using the cor.test function with method = “spearman” and exact = FALSE. Ordinary least-squares trend lines were fitted separately for each treatment comparison and across all comparisons combined. RNA-seq differential expression was determined from gene-level log₂ fold changes, with positive and negative values representing increased and decreased expression, respectively. For m⁶A analysis, significant methylation events were classified according to the reported regulation and fold-change values as hypermethylated or hypomethylated, and transcript-level results were collapsed by gene symbol for gene-level integration. Cd-induced m⁶A reversal was defined as a significant methylation change in Cd versus Vehicle followed by a significant change in the opposite direction in Cd+Akk vs. Cd. AS events were evaluated using event-level *p* values and inclusion-level differences (ΔPSI; IncLevelDifference) and categorized as skipped exon (SE), retained intron (RI), alternative 3′ splice site (A3SS), alternative 5′ splice site (A5SS), or mutually exclusive exon (MXE). An AS event was considered reversed only when the same genomic splicing event was significant in both Cd vs. Vehicle and Cd+Akk vs. Cd and the ΔPSI changed sign between comparisons. Integration across omics layers was performed by gene symbol to identify genes exhibiting concurrent differential expression, m⁶A remodeling, and/or AS regulation, and overlapping genes were further evaluated for concordant, discordant, normalized, or reversed responses following Akk co-treatment. Counts and percentages of overlapping genes were calculated relative to the number of unique genes significantly altered within each m⁶A or AS comparison.

### Data visualization

The distribution density plots were generated using the ggplot2 R package. The Venn diagrams were plotted using the ggVennDiagram R package. Heatmaps were generated from the effect-size estimates for each molecular dataset without z-score transformation. For m⁶A epitranscriptomic data, reported fold-change values were log₂-transformed, with negative and positive values representing hypomethylation and hypermethylation, respectively; genes showing significant Cd-induced changes that were reversed in the Cd+Akk versus Cd comparison were displayed in paired columns. For the integrated alternative-splicing/RNA-seq heatmap, RNA-seq log₂ fold changes were plotted for Cd vs. Vehicle and Cd+Akk vs Cd, while genes were grouped according to the type of significantly reversed AS event: A3SS, A5SS, MXE, RI, or SE. For genes containing multiple reversed AS events, a representative event with the strongest joint statistical support across the two comparisons was used for grouping. Rows in the m⁶A heat maps were grouped into biologically relevant functional/pathway categories based on gene annotation and hippocampal relevance; therefore, these groupings represent functional annotation rather than unsupervised hierarchical clustering. A diverging blue-white-red color scale centered at zero was used, with blue indicating negative and red indicating positive log₂ fold changes. Numerical log₂ fold-change values were displayed within each heat-map cell, and *p* values were presented in separate adjacent columns; an asterisk indicated nominal significance at *p* < 0.05. Pathway-enrichment results were visualized using the Enrichr Appyter Enrichment Analysis Visualizer. Enrichment was treated as exploratory and terms are reported at nominal p unless stated otherwise, consistent with the nominal threshold used throughout the multi-omics integration; where terms also survived Benjamini-Hochberg correction across the ontology, the adjusted P is given.

## RESULTS

Across the hippocampal transcriptome, Cd exposure produced a global shift toward decreased mRNA abundance, accompanied by a biphasic pattern of m⁶A eptranscriptomic changes and alternative-splicing (AS) responses (**Fig. 2A**). *A. muciniphila* (Akk) alone similarly shifted mRNA expression downward and produced a pronounced global decrease in m⁶A, while showing relatively enhanced alternative splicing. In the intervention comparison (Cd+Akk vs Cd), *A. muciniphila* moderately attenuated the Cd-associated decrease in mRNA expression and shifted the m⁶A distribution away from the Cd-induced biphasic pattern toward predominantly hypomethylation, whereas the overall AS distribution was largely unchanged, suggesting little global intervention effect on alternative splicing.

**Figure 2.**
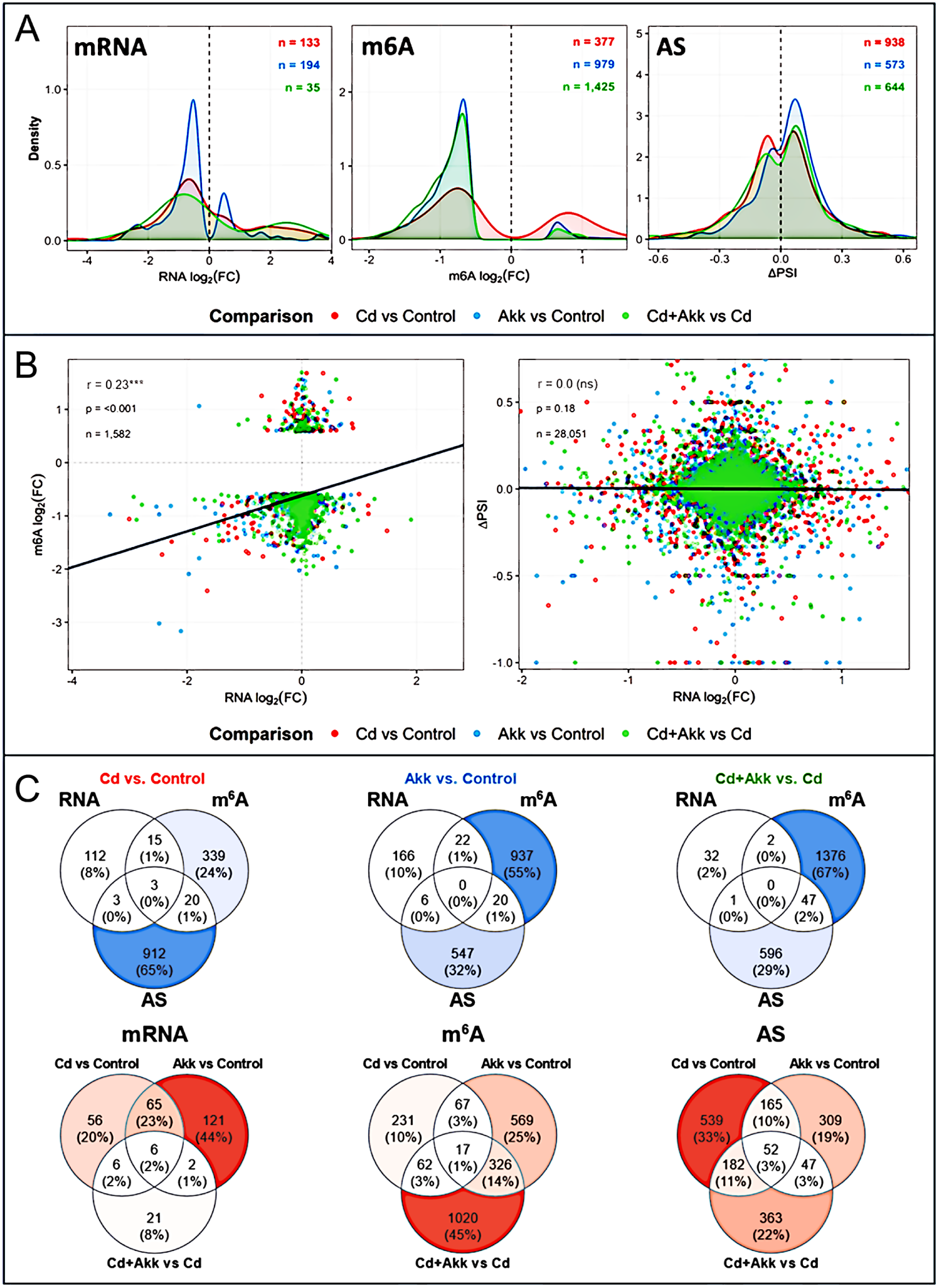
Cd exposure and A. muciniphila differentially affect mRNA expression, m⁶A methylation, and alternative splicing with limited concordance across molecular layers. **A.** Density distributions of significant effect sizes across three molecular layers. Left: RNA log₂ fold changes for differentially expressed genes (Cuffdiff FDR-BH < 0.05; Cd vs. Control n = 133, Akk vs. Control n = 194, Cd+Akk vs. Cd n = 35). Center: m⁶A log₂ fold changes for differentially methylated transcripts (*p* < 0.05; Cd vs. Control n = 377, Akk vs. Control n = 979, Cd+Akk vs. Cd n = 1,425). Right: mean inclusion level differences (ΔPSI) for alternatively spliced genes (rMATS FDR ≤ 0.05; Cd vs. Control n = 938, Akk vs. Control n = 573, Cd+Akk vs. Cd n = 644). Dashed vertical lines indicate zero effect. **B.** Pairwise Spearman rank correlations between omics layer effect sizes across the full tested transcriptome. Each panel shows one pairwise relationship: RNA log₂ fold change versus m⁶A log₂ fold change (left), RNA log₂ fold change versus ΔPSI (centre), and m⁶A log₂ fold change versus ΔPSI (right). Points are coloured by treatment comparison (red: Cd vs. Control, blue: Akk vs. Control, green: Cd+Akk vs. Cd). Per-group linear trend lines and an overall trend line (black) are overlaid without confidence intervals. Spearman ρ values and n per group are annotated in the upper left of each panel. Significance: ns, not significant; \**p* < 0.05; \*\**p* < 0.01; \*\*\**p* < 0.001. **C.** Top panel: three-way Venn diagrams showing the number of genes with significant changes in RNA expression (RNA), m⁶A methylation (m⁶A), or alternative splicing (AS) within each treatment comparison. Percentages indicate the proportion of all detected significant genes in that comparison. Bottom panel: three-way Venn diagrams showing treatment-group overlap for each omics layer independently, illustrating which significant findings are shared across comparisons. Red color intensity reflects the number of genes in each intersection region.

Integration of effect sizes across the three pairwise comparisons showed that steady-state mRNA abundance was largely uncoupled from both m⁶A modification and alternative splicing (**Fig. 2B**). RNA-seq log₂ fold changes exhibited only a weak positive association with m⁶A log₂ fold changes (r = 0.23, *p* < 0.001; n = 1,582), indicating limited concordance between transcript abundance and RNA methylation. In contrast, RNA-seq log₂ fold changes showed essentially no association with alternative-splicing changes measured by ΔPSI (r = 0.00, *p* = 0.18; n = 28,051). Thus, although a modest relationship between mRNA expression and m⁶A was detectable, the overall pattern indicates that changes in total mRNA abundance explain relatively little of the variation in either m⁶A remodeling or splice-isoform regulation, suggesting predominantly post-transcriptional effects of Cd exposure and *A. muciniphila* intervention.

The top three Venn diagrams unveiled the unique and overlapping features (mRNA expression, m⁶A, and AS) among the Cd effect (Cd vs Control), Akk effect (Akk vs Control), as well as the Akk intervention effect against Cd (Cd+Akk vs Cd), as shown in **Fig. 2C**. Regarding the Cd effect (Cd vs. Control), most changes occurred at the post-transcriptional level, with 912 AS-only events (65%) and 339 m⁶A-only genes (24%), whereas only 112 genes (8%) were unique to mRNA expression changes. The overlap among the three features was minimal, with only 3 genes/events shared across mRNA, m⁶A, and AS. Regarding the Akk effect (Akk vs. Control), m⁶A remodeling predominated, with 937 m⁶A only genes (55%), followed by 547 AS-only events (32%) and 166 mRNA-only genes (10%), with no genes shared across all the three features. The strongest m⁶A dominance was observed for the Akk intervention effect against Cd (Cd+Akk vs. Cd), where 1,376 genes (67%) were uniquely altered at the m⁶A level and 596 (29%) uniquely at the AS level, compared with only 32 mRNA-only genes (2%). Together, these data indicate that both Cd exposure and Akk treatment exert much broader effects on RNA methylation and splice-isoform regulation than on total mRNA abundance, with Akk-mediated changes particularly enriched at the m⁶A level from both Akk alone and the Akk remediation groups.

The bottom three Venn diagrams in Fig. 2B compared the degree of overlap among the three types of exposures (Cd effect: Cd vs. Control, Akk effect: Akk vs. Control, and Akk intervention effect against Cd: Cd+Akk vs. Cd) within each molecular feature (mRNA, m⁶A, and AS). For mRNA expression, effects were largely exposure-specific: 56 genes were unique to Cd effect, 121 to Akk effect, and 21 to Akk intervention against Cd, while only 6 genes (2%) were common to all three exposures. In contrast, the m⁶A dataset showed a particularly strong Akk intervention-specific response, with 1,020 genes (45%) unique to the Cd+Akk vs Cd comparison, along with 326 genes (14%) shared between Akk alone and Akk intervention, consistent with a substantial Akk-associated m⁶A remodeling signature. Alternative splicing (AS) changes were more broadly distributed, with 539 events (33%) unique to Cd effect, 309 (19%) unique to Akk alone, and 363 (22%) unique to Akk intervention. Notably, 182 AS events (11%) were shared between Cd and Akk intervention, while 52 events (3%) were common to all three comparisons. Overall, across the molecular features, responses were largely exposure-specific, with Akk intervention against Cd producing especially strong m⁶A remodeling, while mRNA changes showed limited overlap and alternative splicing responses were more broadly shared across exposures.

To examine the effect of *A. muciniphila* intervention on the Cd-induced hippocampal transcriptomic response, pathway analyses were conducted on genes that overlapped between the Cd vs. Control and Cd + Akk vs. Cd comparisons (Fig. 3). Interestingly, all overlapping genes exhibited a reversal pattern, such that genes upregulated by Cd were downregulated by *A. muciniphila* co-exposure, whereas genes downregulated by Cd were upregulated by *A. muciniphila*, indicating a broad remediation of Cd-induced transcriptional dysregulation. The complete reversal of the Cd-responsive genes by *A. muciniphila* was associated with biological processes highly relevant to hippocampal function and neuronal resilience (Fig. 3A). Neuropeptide signaling, the most significantly enriched Gene Ontology process, is important for neurological functions because neuropeptides are potent modulators of neuronal ion channels, synaptic transmission, network plasticity, and behavior (Smith et al., 2020). Enrichment of negative regulation of membrane depolarization further points to restoration of mechanisms controlling neuronal excitability, whereas negative regulation of mitochondrial depolarization suggests normalization of mitochondrial membrane-potential control; synaptic mitochondria are essential for ATP production, Ca²⁺ buffering, neurotransmitter release, and maintenance of synaptic activity, and mitochondrial dysfunction is strongly linked to synaptic failure and neurodegeneration (Ly and Verstreken, 2006; Lee et al., 2018). Regulation of cell-adhesion molecule production is also neurologically important because neural cell-adhesion molecules such as NCAM and L1 participate directly in hippocampal long-term potentiation and learning-related synaptic remodeling (Luthi et al., 1994; Bukalo et al., 2004). The enrichment of cholesterol import and lipoprotein-remodeling pathways further implicates restoration of neuronal lipid homeostasis, because cholesterol availability is required for synaptic membrane organization and hippocampal plasticity, and experimental depletion of hippocampal cholesterol impairs LTP and memory (Guo et al., 2021). Finally, the enrichment of protein oxidation and protein processing links the reversal signature to redox damage and proteostasis, processes to which postmitotic neurons are particularly vulnerable and whose disruption is a common feature of neurodegenerative disease (Tseng et al., 2023).

**Figure 3.**
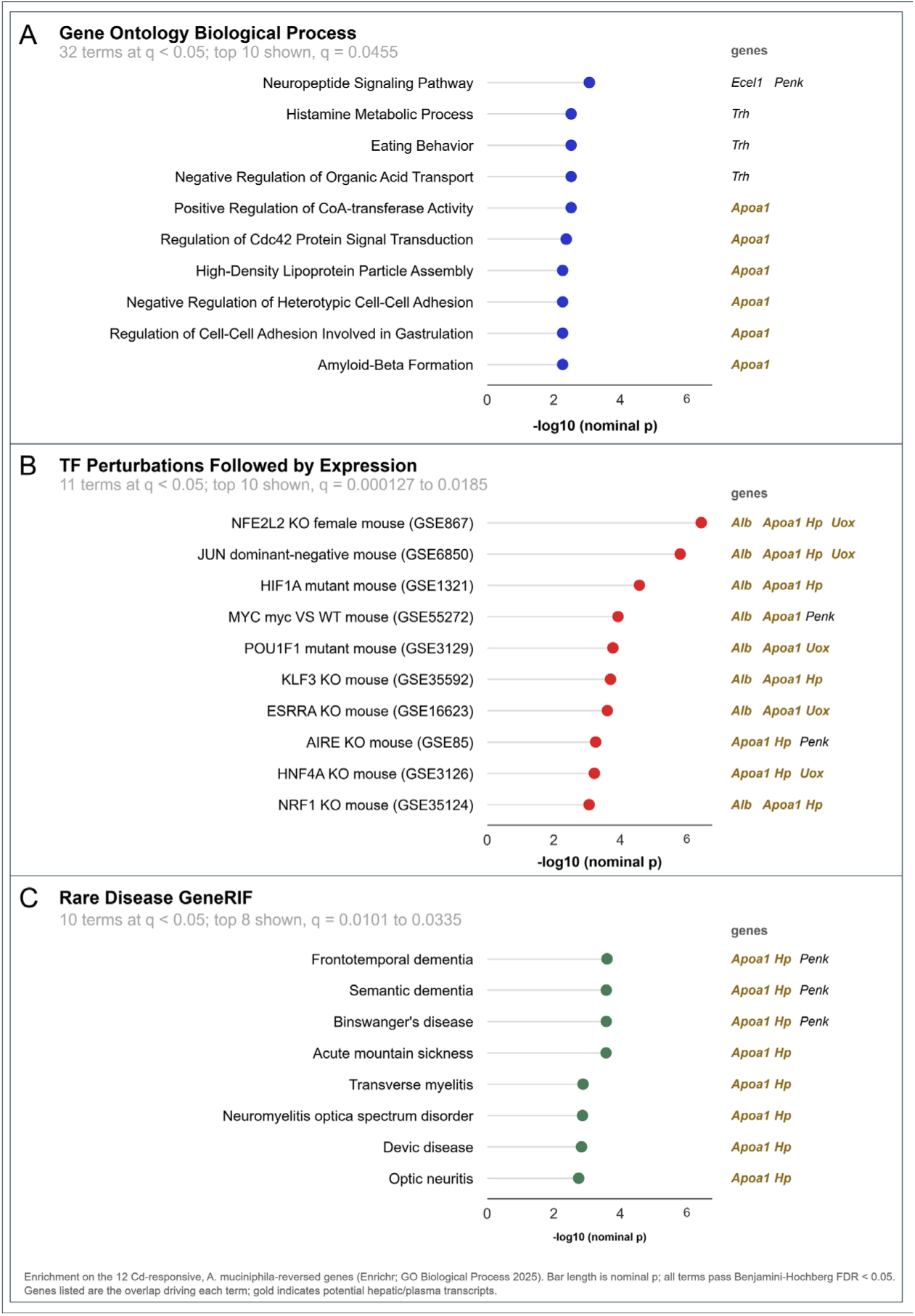
Functional enrichment analysis of the Cd-responsive, *A. muciniphila*-reversed transcripts. Messenger RNAs significantly altered by Cd versus Vehicle and altered in the opposite direction by Cd+Akk versus Cd (Benjamini–Hochberg FDR < 0.05; 12 genes, Figure 4A) were subjected to enrichment analysis across three libraries. **A.** Gene Ontology Biological Process; 32 terms reached FDR < 0.05 and the ten most significant are shown. **B.** TF Perturbations Followed by Expression; 11 terms reached FDR < 0.05, one signature per transcription factor, ten shown. **C.** Rare Disease GeneRIF; 10 terms reached FDR < 0.05, eight shown. Point position is the nominal −log10 p; all terms shown pass Benjamini–Hochberg FDR <0.05. Genes beside each term are the overlap driving it, with hepatic and plasma transcripts (Alb, Apoa1, Hp, Mup20, Serpina1e, Uox) in gold. Half of the input list is hepatic or plasma in origin, and those transcripts carry the enrichment: across all three libraries the significant terms rest on the same three or four genes, and the NFE2L2 and JUN signatures in B share an identical overlap. The panels therefore describe the composition of the gene list more than they identify a transcription-factor or disease program, and the four neuronal genes in the set (Ecel1, Npas3, Trh, Penk) contribute to only one term.

Most notably, transcription-factor perturbation analysis revealed a striking NFE2L2/Nrf2-centered signature, with Nrf2 perturbations accounting for four of the ten highest-ranking signatures due to *A. muciniphila* intervention (Fig. 3B). Nrf2 is the master cytoprotective sensor against oxidative stress and electrophiles (Kobayashi et al., 2006; Yamamoto et al., 2018). This predicted pathway is important for neutralizing Cd neurotoxicity, because oxidative stress is a major mechanism of Cd-induced CNS injury (Branca et al., 2020), and experimental Cd exposure has directly implicated Nrf2 in protection against Cd-induced synaptic damage, microglial activation, and learning/memory impairment (Li et al., 2025). Nrf2 is also highly relevant to brain resilience: NRF2-dependent antioxidant programs in astrocytes protect neighboring neurons against oxidative injury, in part by supporting glutathione-dependent neuronal defense (Vargas and Johnson, 2009; Jimenez-Blasco et al., 2015). Other enriched regulators for Akk intervention include JUN/c-JUN, a stress-responsive component of AP-1 signaling, participates directly in hippocampal plasticity, with c-JUN phosphorylation required for normal CA3-CA1 long-term potentiation (Seo et al., 2012); HIF1A, which couples oxygen/redox state to cellular metabolism and can profoundly affect hippocampal synaptic plasticity and memory when dysregulated (Arias-Cavieres et al., 2020); as well as estrogen-related receptor alpha (ESRRA/ERRα), which works with peroxisome proliferator-activated receptor gamma coactivator 1-alpha (PGC-1α) to control mitochondrial and synaptic gene programs in neurons, including in the hippocampus (McMeekin et al., 2021). Together, *A. muciniphila*-mediated protection against Cd-induced memory deficit (Wang et al., 2026a) is associated with a reversal of Cd-induced hippocampal transcriptional changes involving predicted interconnected pathways related to Nrf2-centered redox defense, mitochondrial bioenergetics, neuronal excitability, synaptic plasticity, lipid homeostasis, and proteostasis. These coordinated transcriptional patterns may represent a molecular signature consistent with potential remediation of Cd-induced hippocampal perturbations.

At the gene level, RNA-seq analysis identified 12 hippocampal genes of which Cd-induced expression changes were reversed in direction by *A. muciniphila* co-exposure (Fig. 4 and Table 1). Four genes associated with neuronal signaling, plasticity, and adult neurogenesis were markedly down-regulated by Cd and up-regulated following Akk co-treatment: neuronal PAS domain protein 3 (Npas3) (Cd log₂FC = –2.79; Cd+Akk vs Cd = +1.26), thyrotropin-releasing hormone (Trh) (–2.27; +2.27), proenkephalin (Penk) (–1.45; +0.56), and endothelin-converting enzyme-like 1 (Ecel1) (–1.28; +1.24). In contrast, Kruppel-like factor 8 (Klf8) and pleckstrin and Sec7 domain-containing protein 4 (Psd4), grouped with neuronal transcriptional regulation and membrane trafficking, were induced by Cd (+2.27 and +2.96, respectively) and decreased with Akk co-treatment (–2.25 and –4.65). Thus, the neuronal-related genes showed a coordinated bi-directional reversal pattern, with *A. muciniphila* opposing both Cd-associated suppression of genes linked to neurogenesis/neuromodulation and Cd-associated induction of genes involved in transcriptional and membrane-remodeling responses. A second group of Cd-responsive genes associated with neurovascular/inflammatory, lipid-transport, metabolic, and redox-stress functions was uniformly up-regulated by Cd and down-regulated following Akk co-treatment. These included albumin (Alb) (+2.67 ◊ –2.19), apolipoprotein A-I (Apoa1) (+2.55 ◊ – 2.55), haptoglobin (Hp) (+2.05 ◊ –2.38), and serine peptidase inhibitor, clade A, member 1e (Serpina1e) (+2.56 ◊ –2.67), consistent with reversal of a Cd-associated vascular/acute-phase and lipid-response transcriptional signature. In parallel, urate oxidase (Uox) (+2.69 ◊ –3.03) and major urinary protein 20 (Mup20) (+3.44 ◊ –3.02) were grouped with metabolic and stress-response functions and likewise showed complete directional reversal. Collectively, these RNA-seq data indicate that *A. muciniphila* co-treatment opposed the direction of Cd-induced transcriptional changes across all 12 genes, suggesting partial normalization of a coordinated hippocampal expression signature involving neuronal function, neurovascular/inflammatory responses, lipid handling, and cellular stress pathways. This finding provides a potential molecular explanation for the *A. muciniphila*-mediated prevention of Cd-induced memory deficits that we previously observed from the same mouse cohort (Wang et al., 2026a).

**Figure 4.**
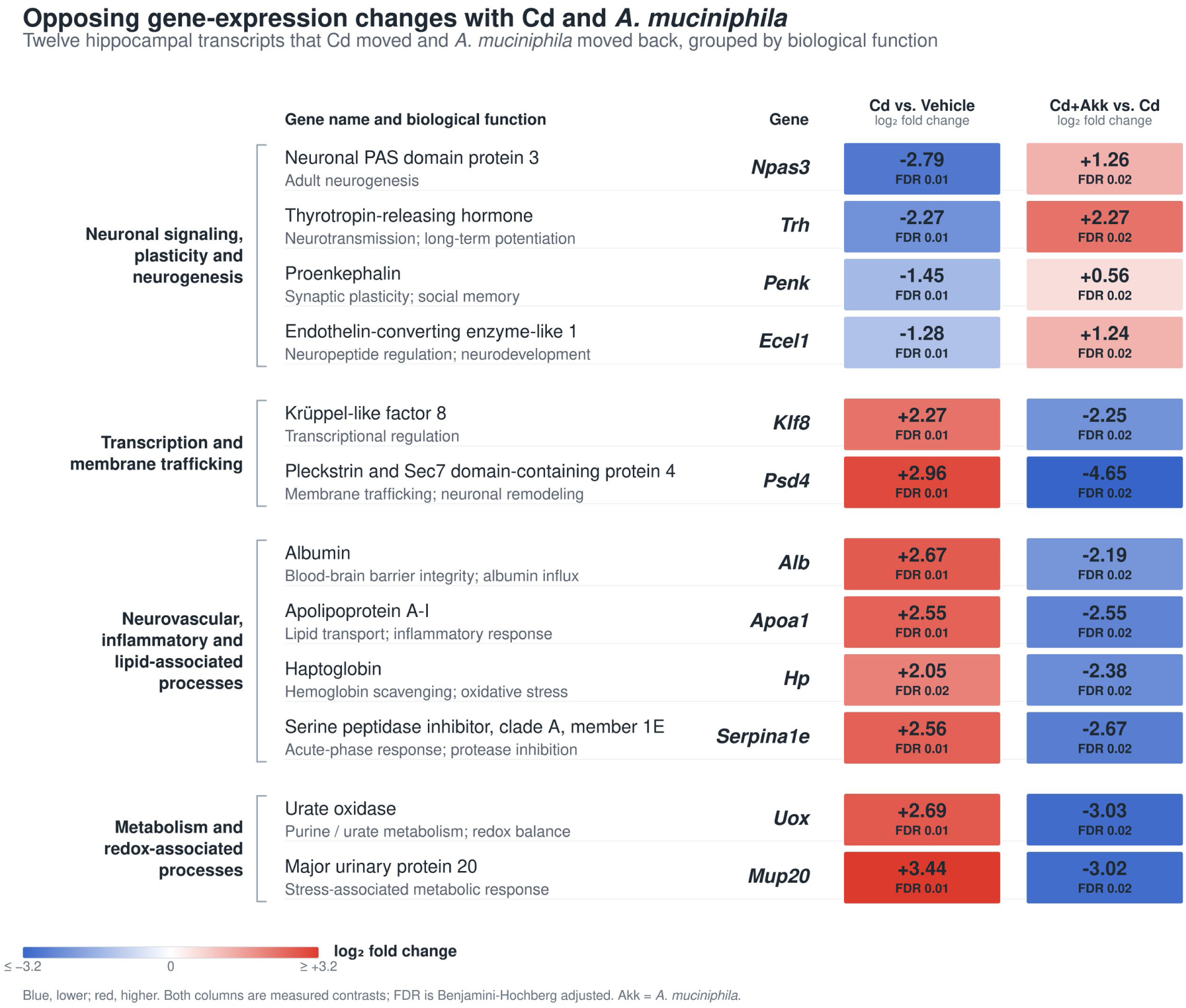
*A. muciniphila* reverses key Cd-responsive hippocampal transcriptional signatures. Heatmap of 12 hippocampal transcripts showing significant directional reversal between Cd vs. Control and Cd+Akk vs. Cd. Cd decreased *Npas3, Trh, Penk,* and *Ecel1*, whereas Akk co-treatment shifted these transcripts in the opposite direction. Cd increased *Hp, Klf8, Apoa1, Serpina1e, Alb, Uox, Psd4,* and *Mup20*, and these changes were similarly opposed by Akk intervention. Values are RNA-seq log₂ fold changes with Benjamini–Hochberg FDR. Red indicates increased and blue indicates decreased expression. Transcripts are grouped by literature-based biological function (PubMed): neuronal signaling, plasticity and neurogenesis; transcription and membrane trafficking; neurovascular, inflammatory and lipid-associated processes; and metabolism and redox-associated processes. Functional groupings are intended to summarize biological relationships rather than represent unsupervised clustering.

**Table 1.** *A. muciniphila* remediated hippocampal mRNAs against Cd exposure.

|  |  |  |  |  |  |
| --- | --- | --- | --- | --- | --- |
| **Gene Symbols** | **Cd over Control**<br>**log<sub>2</sub>(fold change)** | **FDR-BH**<br>**(Cd over Control)** | **Cd+Akk over Cd**<br>**log<sub>2</sub>(fold change)** | **FDR-BH**<br>**(Cd+Akk over Cd)** | **Annotation** |
| **Penk** | -1.45 | 0.01 | 0.56 | 0.02 | Proenkephalin; precursor of endogenous enkephalins involved in opioid neuromodulation. Genetic loss of Penk alters neuronal activation in hippocampal CA1–CA3 during cue-associated behavior, supporting relevance to hippocampal memory/reward circuitry (Gutierrez-Cuesta et al., 2014) |
| **Ecel1** | -1.28 | 0.01 | 1.24 | 0.02 | Endothelin-converting enzyme-like 1; neuronal membrane-bound metalloprotease important for neuronal development and axonal arborization (Matsumoto et al., 2016) |
| **Npas3** | -2.79 | 0.01 | 1.26 | 0.02 | Neuronal PAS domain protein 3; neuronal transcription factor required for normal adult hippocampal neurogenesis. Loss of Npas3 reduces hippocampal FGF receptor signaling and neurogenesis (Pieper et al., 2005) |
| **Hp** | 2.05 | 0.02 | -2.38 | 0.02 | Haptoglobin; hemoglobin-binding acute-phase protein with antioxidant and immunomodulatory functions. In the CNS, Hp regulates macrophage/microglial inflammatory responses (Morimoto et al., 2022) |
| **Klf8** | 2.27 | 0.01 | -2.25 | 0.02 | Krüppel-like factor 8; nuclear transcription factor that is highly expressed in neurons throughout the adult mouse brain, including the hippocampus; involved in cell signaling and proliferation (Dobrivojevic et al., 2015) |
| **Trh** | -2.27 | 0.01 | 2.27 | 0.02 | Thyrotropin-releasing hormone; CNS neuropeptide with neurotransmitter/neuromodulator actions. In hippocampal slices, TRH suppresses depolarization-evoked glutamate and aspartate release, supporting a role in regulation of hippocampal excitability (Nie et al., 2005) |
| **Alb** | 2.67 | 0.01 | -2.19 | 0.02 | Plasma transport protein; hippocampal expression may reflect vascular/barrier or injury-associated responses (Braganza et al., 2012) |
| **Apoa1** | 2.55 | 0.01 | -2.55 | 0.02 | Lipid/cholesterol transport protein relevant to neuronal membrane and lipid homeostasis; Increased circulating ApoA-I preserves learning/memory and cerebrovascular integrity in APP/PS1 mice (Lewis et al., 2010) |
| **Serpina1e** | 2.56 | 0.01 | -2.67 | 0.02 | Serpin-family protein potentially associated with protease and inflammatory regulation, Serpina1e can enhance hippocampal memory, neurogenesis, BDNF expression, and neurite growth (Kim et al., 2026) |
| **Uox** | 2.69 | 0.01 | -3.03 | 0.02 | Urate oxidase; enzyme of uric-acid metabolism, potentially related to redox/metabolic homeostasis, controls uric-acid/urate metabolism and thereby influences systemic and brain redox metabolism. Manipulation of Uox alters brain urate concentrations and neurochemical phenotypes (Chen et al., 2013) |
| **Mup20** | 3.44 | 0.01 | -3.02 | 0.02 | Major urinary protein 20 (darcin); classically a male pheromone protein. Interestingly, exposure to darcin can stimulate adult hippocampal neurogenesis in female mice (Hoffman et al., 2015) |
| **Psd4** | 2.96 | 0.01 | -4.65 | 0.02 | Pleckstrin and Sec7 domain-containing 4; ARF-family guanine nucleotide-exchange factor involved in membrane trafficking/recycling and cytoskeletal-associated signaling. These processes could be relevant to neuronal membrane dynamics (Hongu et al., 2015) |

As shown in **Fig. 5** (left panel), among Cd-induced m⁶A epitranscriptomic alterations in the hippocampus, 32 genes that were significantly hypermethylated by Cd relative to vehicle became significantly hypomethylated following *A. muciniphila* intervention, demonstrating a coordinated reversal of Cd-induced m⁶A regulation. The m⁶A reversed genes represented several functional modules, including Ca²⁺/cyclic-nucleotide and receptor signaling, membrane/cytoskeletal organization, DNA-damage and redox regulation, and immune/metabolic signaling. Consistent with these patterns, pathway analysis (**Fig. 6A**) highlighted processes related primarily to immune regulation and cellular stress/genome maintenance. The strongest enrichment was negative regulation of αβ T-cell activation (*p* = 1.62×10⁻⁴), accompanied by regulation of αβ T-cell proliferation and CD8⁺ T-cell activation, suggesting that Cd-sensitive m⁶A remodeling involves transcripts annotated to immune-modulatory pathways. Additional enriched processes included mitotic spindle localization, response to ionizing radiation, telomere capping and protein localization to telomeres, consistent with regulation of genes involved in cell-cycle organization, DNA-damage responses, and genome stability. Cellular response to histamine and vitamin K metabolism were also enriched. The p values reported for these terms are nominal. After Benjamini-Hochberg correction across the ontology, only negative regulation of αβ T-cell activation remained significant (adjusted P = 0.024; Dapl1 and Tnfrsf14). Overall, these findings suggest that Akk-mediated reversal of Cd-induced m⁶A hypermethylation preferentially affected transcripts associated with immune signaling, cellular stress responses, and genome-maintenance processes.

**Figure 5.**
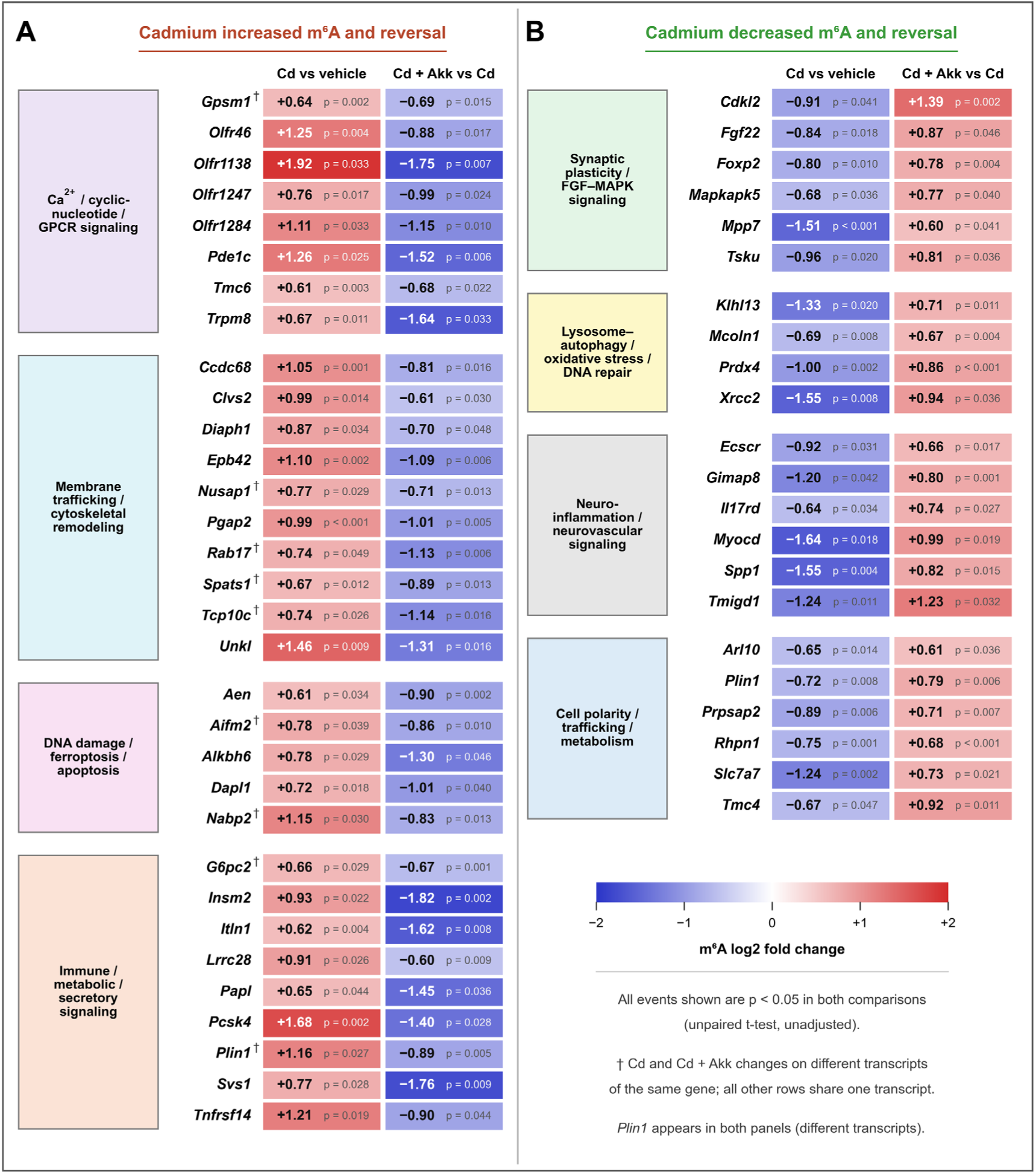
*A. muciniphila* reverses Cd-induced hippocampal m⁶A epitranscriptomic remodeling. Heatmaps of Cd-induced m⁶A alterations that were significantly reversed by Akk co-treatment. (A) Thirty-two gene-level events showed Cd-induced hypermethylation followed by hypomethylation in Cd+Akk vs. Cd, encompassing Ca²⁺/cyclic-nucleotide/GPCR signaling, membrane trafficking and cytoskeletal remodeling, DNA-damage/ferroptosis/apoptosis, and immune/metabolic/secretory signaling. (B) Twenty-two gene-level events showed the reciprocal pattern of Cd-induced hypomethylation followed by hypermethylation, involving synaptic plasticity/FGF–MAPK signaling, lysosome–autophagy/oxidative-stress/DNA-repair pathways, neuroinflammatory/neurovascular signaling, and cellular trafficking/metabolism. Because *Plin1* was represented in both directional categories, these reversal events correspond to 53 unique genes. Values are m⁶A log₂ fold changes (red, hypermethylation; blue, hypomethylation) with unadjusted t-test p values; every event shown is p < 0.05 in both comparisons. † Cd and Cd+Akk changes detected on different transcripts of the same gene.

**Figure 6.**
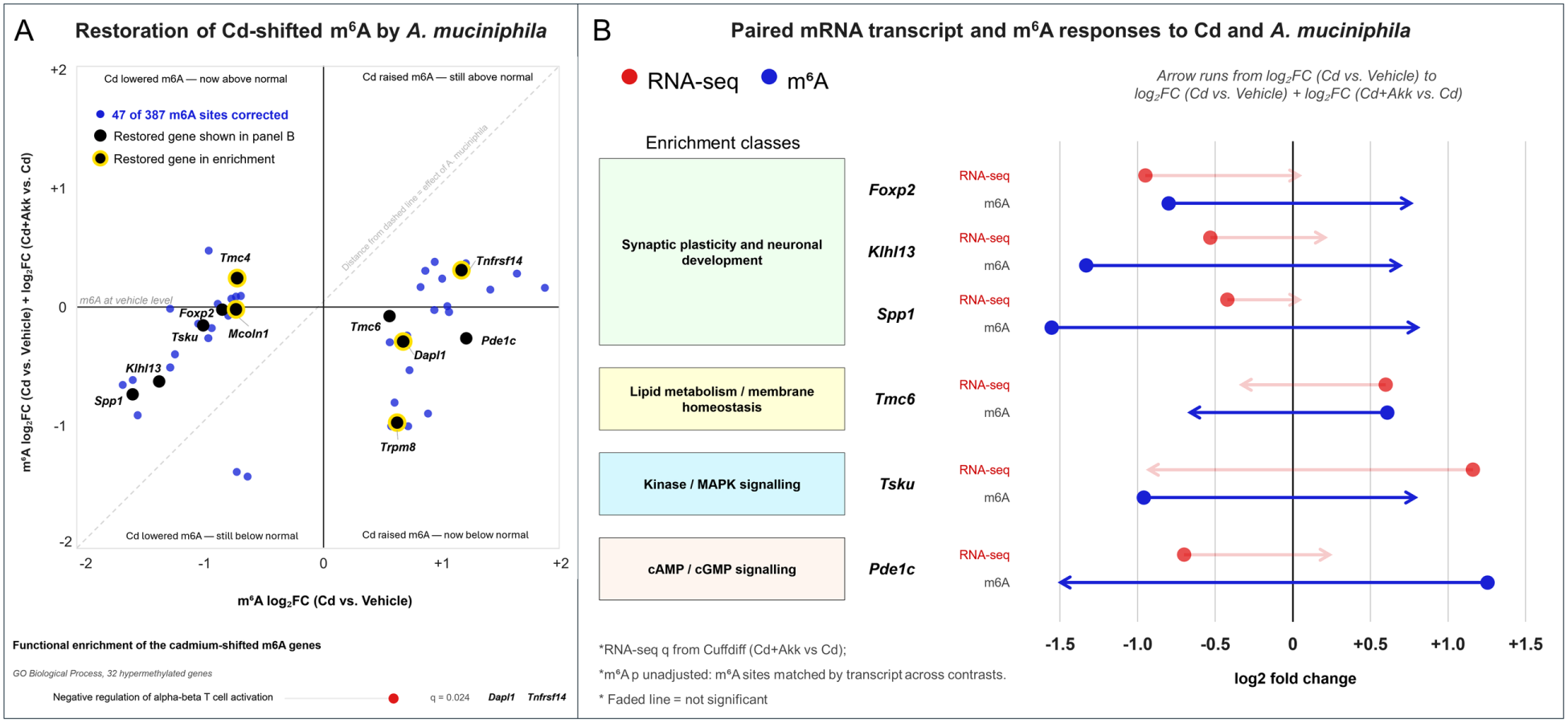
*A. muciniphila* reverses Cd-shifted m⁶A methylation. **A.** Cd significantly shifted m⁶A at 387 transcripts; the 47 that the Cd+Akk vs. Cd comparison also called significant are plotted as the measured Cd effect (x-axis) against the effect remaining under co-treatment relative to vehicle (y-axis), the latter calculated as the sum of the two measured comparisons, because a Cd+Akk vs. Vehicle contrast was not generated for the m⁶A array. The dashed diagonal marks no effect of A. muciniphila, so distance below it is the size of the A. muciniphila effect; the horizontal axis marks return to the vehicle level. m⁶A sites were matched by transcript identifier across the two comparisons. Black points, genes also shown in panel B; yellow rings, genes carrying an enriched term. Below, functional enrichment of the Cd-shifted m⁶A genes: negative regulation of alpha-beta T cell activation (GO Biological Process, 32 hypermethylated genes, adjusted P = 0.024), the only term in either discordant set to survive Benjamini–Hochberg correction. Ion-channel terms in the Molecular Function ontology, driven by Tmc4, Tmc6, Mcoln1 and Trpm8, are enriched at nominal p but fall at adjusted P ≥ 0.051 and are therefore shown on the plot, but not reported as significant. **B.** Six genes with both m⁶A and transcript measurements, grouped by function. Each gene has two tracks; the arrow runs from the Cd effect to the effect under co-treatment, in log₂ fold change. Faded arrows are not significant. Cd moved m⁶A and transcript abundance in the same direction for four of the six genes and in opposite directions for Tsku and Pde1c.

As shown in **Fig. 5** (right panel), among Cd-induced m⁶A epitranscriptomic alterations in the hippocampus, 22 genes that were significantly hypomethylated by Cd relative to vehicle became significantly hypermethylated following *A. muciniphila* intervention. The m⁶A reversed genes represented several functional modules, including synaptic plasticity/FGF-MAPK signaling, lysosome–autophagy/oxidative stress/DNA repair, neuroinflammatory/neurovascular signaling, and cell polarity/trafficking/metabolism. Consistent with these patterns, pathway analysis (**Fig. 6A**) highlighted processes related primarily to cell differentiation, nucleotide/ribose-phosphate metabolism, and transforming growth factor-β1 (TGF-β1) regulation. The strongest enrichment was negative regulation of cell differentiation (*p* = 1.85×10⁻³), followed by 5-phosphoribose-1-diphosphate biosynthetic and metabolic processes (P= 5.49×10⁻³), negative regulation of TGF-β1 production (*p* = 5.49×10⁻³), and ribose-phosphate biosynthetic process (*p* = 6.58×10⁻³), indicating involvement of pathways related to cellular differentiation and nucleotide metabolism. These terms are reported at nominal p; none reached Benjamini-Hochberg significance across the ontology, and several derive from a single gene, Myocd. Additional enriched terms included cardiac vascular smooth muscle cell differentiation, negative regulation of skeletal muscle cell differentiation, positive regulation of cardiac muscle cell differentiation, response to testosterone, and cardiac muscle cell myoblast differentiation. Although several of these GO terms carry muscle-related annotations, in the hippocampal context they likely reflect broader cell-differentiation, cytoskeletal, and neurovascular regulatory pathways. Overall, these findings suggest that *A. muciniphila*-mediated reversal of Cd-induced m⁶A hypomethylation preferentially affected transcripts associated with neuronal plasticity, stress-response and genome-maintenance pathways, cell differentiation, and metabolic/nucleotide biosynthetic processes.

Integrative analysis of RNA-Seq and m⁶A epitranscriptomics identified six hippocampal transcripts in which Cd-induced m⁶A alterations were reversed by *A. muciniphila*, with corresponding reversal of mRNA expression in at least one of the paired-wise comparisons (Cd vs. Control, or Cd+Akk vs Cd) (**Fig. 6B**). Genes involved in synaptic plasticity and neuronal development, namely Forkhead box P2 (Foxp2), kelch-like family member 13 (Klhl13), and secreted phosphoprotein 1 (Spp1) were significantly downregulated by Cd at the mRNA level (log_2_ fold change = –0.951, –0.531, and –0.422, respectively) and simultaneously m⁶A-hypomethylated (log_2_ fold change = –0.799, –1.331, and –1.553). Cd+Akk significantly reversed their m⁶A status to hypermethylation (log_2_ fold change = +0.779, +0.710, and +0.823), while their mRNA levels had an increasing trend but not statistically significant. To note, Foxp2 is an established regulator of neural-circuit development, learning, and synaptic plasticity, with Foxp2 perturbation altering neuronal plasticity and skill learning (Groszer et al., 2008; Schreiweis et al., 2014). Klh13 functions as a CUL3-associated ubiquitin-ligase adaptor involved in cell-cycle regulation, and pathogenic Klh13 variants have recently been linked to neurodevelopmental disorders and genomic instability (Akhter et al., 2026). Spp1/osteopontin has direct hippocampal relevance: perivascular Spp1 induces microglial phagocytic states and promotes microglial synaptic engulfment in mouse models of Alzheimer’s disease (De Schepper et al., 2023). In contrast, transmembrane channel-like 6 (Tmc6) was increased by Cd at both the mRNA (+0.599) and m⁶A (+0.608) levels; Cd+Akk significantly reversed its m⁶A modification to hypomethylation (–0.682), while its mRNA expression had a decreasing trend although not statistically significant. To note, Tmc6 has been shown to regulate neuronal membrane excitability through modulation of M-type potassium channels in primary sensory neurons (An et al., 2024). Interestingly, inverse RNA-m⁶A relationships were also observed for tsukushi, small leucine-rich proteoglycan (Tsku) and phosphodiesterase 1C (Pde1c). Cd markedly increased Tsku mRNA (log_2_ fold change = +1.161) while decreasing its m⁶A modification (log_2_ fold change = –0.959); Cd+Akk reversed both responses, significantly reducing Tsku mRNA (–0.947) and increasing m⁶A (log_2_ fold change = +0.808). Tsku is directly implicated in hippocampal biology, where it regulates dentate-gyrus neural stem-cell maintenance, proliferation, and terminal differentiation (Ahmad et al., 2020). Conversely, Cd decreased Pde1c mRNA (log_2_ fold change = –0.700) while increasing its m⁶A modification (log_2_ fold change = +1.255), whereas Cd+Akk strongly reversed the m⁶A change (log_2_ fold change = –1.515), with an increasing mRNA trend (although not statistically significant). To note, Pde1c is a Ca²⁺/calmodulin-dependent phosphodiesterase that hydrolyzes both cAMP and cGMP and is expressed in multiple neuronal regions of the brain (Yan et al., 1996), linking this reversal to cyclic-nucleotide signaling. To determine whether the m⁶A sites reversed by A. muciniphila returned toward the vehicle baseline rather than simply moving in the opposite direction, we compared the Cd effect at each site with the effect remaining under co-treatment, both relative to vehicle (**Fig. 6A**). Cd significantly shifted m⁶A at 387 transcripts. Forty-seven of these were also called significant in the Cd+Akk vs. Cd comparison, and 45 of those 47 (96%) moved in the opposing direction; 22 returned to within 0.25 log₂ units of the vehicle level, while six were displaced further from it. Sites were matched by transcript identifier across the two comparisons, since several genes carry more than one m⁶A site and matching by gene symbol pairs different transcripts. Forty-five of these transcripts fall within the 53 genes identified by gene-level matching; for the remaining eight of those 53, the site shifted by Cd and the site shifted by co-treatment are different transcripts of the same gene. Functional enrichment of the Cd-shifted m⁶A genes identified negative regulation of alpha-beta T cell activation among the 32 hypermethylated genes (adjusted P = 0.024, Dapl1 and Tnfrsf14), the only term in either discordant set to survive Benjamini–Hochberg correction. Ion-channel terms across the 53 discordant genes, driven by Tmc4, Tmc6, Mcoln1 and Trpm8, were enriched at nominal p but fell at adjusted P ≥ 0.051.

Overall, integrated RNA-seq and m⁶A analyses indicate that *A. muciniphila* counteracts Cd-induced hippocampal dysregulation accompanied by restoring transcript-specific m⁶A states of a subset of genes related to neurological pathways and signaling transduction, however, because the vast majority of m⁶A reversals occurred without corresponding changes in total mRNA abundance, these effects likely reflect predominantly post-transcriptional mechanisms such as RNA stability, splicing, localization, and especially translational control rather than changes in total transcript abundance, which needs to be addressed in future directions.

Table 2 summarizes the *A. muciniphila*-mediated reversal of Cd-induced alternative-splicing events by directly comparing Cd+Akk vs Cd against Cd vs Vehicle. Across all AS categories, 1,267 individual splicing events involving 1,052 unique genes showed significant reversal in direction. Skipped exon (SE) events represented the largest class, with 807 reversed events affecting 705 genes, followed by alternative 3′ splice site (A3SS) events (156 events, 148 genes), retained intron (RI) events (135 events, 130 genes), alternative 5′ splice site (A5SS) events (89 events, 86 genes), and mutually exclusive exon (MXE) events (80 events, 76 genes). Integration with RNA-seq showed that only a small proportion of genes with reversed AS events also exhibited significant changes in total mRNA abundance: 13 A3SS genes (8.8%), 10 A5SS genes (11.6%), 4 MXE genes (5.3%), 10 RI genes (7.7%), and 36 SE genes (5.1%) overlapped with differentially expressed genes. Overall, only 63 of the 1,052 AS-reversal genes (6.0%) were also DEGs, indicating that the vast majority of Akk-mediated reversal of Cd-induced splicing dysregulation occurred without corresponding changes in steady-state mRNA abundance. Pathway enrichment analysis of the 63 genes revealed enrichment for neuronal excitability, synaptic transmission, ion transport, and neurodevelopmental regulation. The most significantly enriched processes were modulation of excitatory postsynaptic potential (*p* = 1.38×10⁻⁴) and negative regulation of excitatory postsynaptic potential (*p* = 1.45×10⁻⁴), followed by regulation of metal ion transport (*p* = 2.14×10⁻⁴), regulation of neurotransmitter secretion (*p* = 3.85×10⁻⁴), and negative regulation of sodium ion transport (*p* = 4.32×10⁻⁴). Additional enriched processes included positive regulation of Wnt signaling, modulation of chemical synaptic transmission, regulation of cell projection organization, phosphorylation, and regulation of neuron differentiation. As shown in Supplementary Figure S1, only three genes, namely calcium/calmodulin-dependent protein kinase II delta (Camk2d), neuroglobin (Ngb), and procollagen C-endopeptidase enhancer (Pcolce), showed significant opposite-direction RNA-seq changes in both comparisons, consistent with coordinated reversal at both the splicing and transcript-abundance levels. These findings further support a predominantly post-transcriptional mechanism of Akk action, with extensive restoration of splice-isoform regulation that would not be captured by gene-level RNA-seq analysis alone. To identify the events most likely to represent genuine restoration rather than the mirror-image effects expected from a shared Cd reference group, we applied a more stringent definition requiring FDR-corrected significance in both comparisons, a Cd effect of at least 10% PSI, and return to the vehicle baseline in the fourth contrast (Cd+Akk vs. Vehicle). Ninety-nine events in 92 unique genes met all four criteria (Table 2, right-hand columns), again with skipped exons as the largest class (56 events, 51 genes). These high-confidence restoration events are shown in Figure 7 and listed individually in Supplementary Table S1. Functional enrichment of the 92 restored genes identified three terms surviving Benjamini-Hochberg correction (Fig. 7B): mRNA 3’-UTR AU-rich region binding (adjusted P = 2.3 x 10-5; Arid5a, Cpeb1, Cpeb2, Exosc8), calcium-induced calcium release activity (adjusted P = 0.017; Ryr1, Ryr3) and cellular response to insulin stimulus (adjusted P = 0.036). The 32 restored genes with the largest Cd effects are shown gene by gene in Figure 7C, with the percentage of the Cd effect reversed given for each. Nine of the 92 genes (9.8%) were also differentially expressed by RNA-seq, a proportion that remains close to the rate expected by chance given the number of genes tested in each dataset (approximately 6.6%), reinforcing the conclusion that splice-isoform restoration occurs largely independently of transcript abundance. Across all 7,751 genes carrying a reversed splicing event for which RNA-seq data were available, the median absolute transcript-level change under Cd was 0.10 log₂ units and only 53 genes (0.7%) were differentially expressed at q < 0.05 (Supplementary Figure S2).

**Figure 7.**
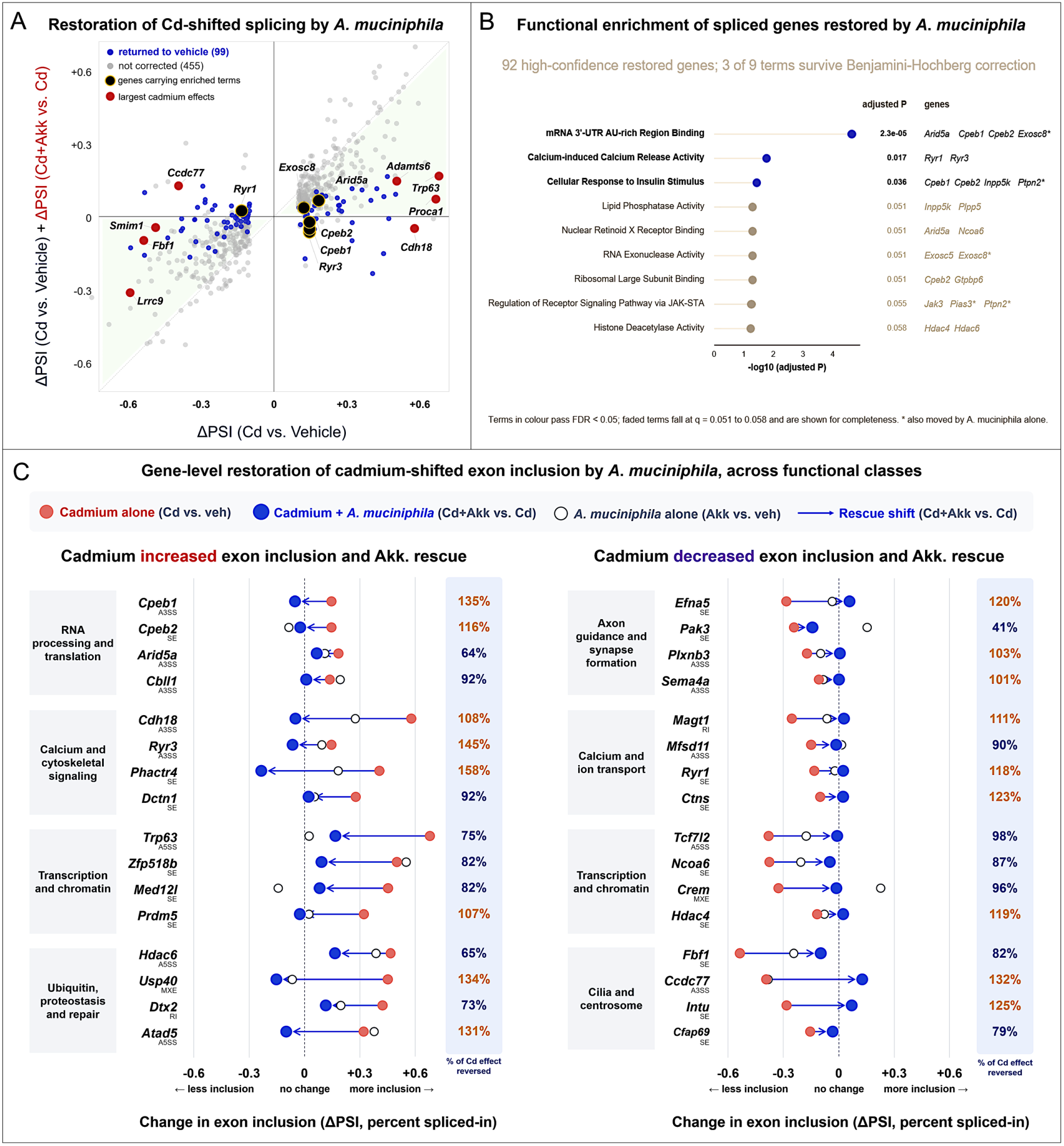
High-confidence restoration of Cd-disrupted alternative splicing by *A. muciniphila*. **A.** All 554 splicing decisions significantly altered by Cd (FDR < 0.05, |ΔPSI| ≥ 0.10), plotted as the measured Cd effect against the measured effect remaining under co-treatment relative to vehicle (Cd+Akk vs. Vehicle). Points on the diagonal were not corrected; points on the horizontal returned to the vehicle level. Blue, the 99 decisions in 92 genes meeting the high-confidence restoration criterion — opposite direction at FDR < 0.05 in both comparisons, a Cd effect of at least 10% PSI, and no residual departure from the vehicle baseline (Cd+Akk vs. Vehicle, FDR ≥ 0.05). Grey, decisions not corrected. **B.** Functional enrichment of the 92 restored genes. Three of nine terms survive Benjamini–Hochberg correction: mRNA 3′-UTR AU-rich region binding (adjusted P = 2.3 × 10−5), calcium-induced calcium release activity (adjusted P = 0.017) and cellular response to insulin stimulus (adjusted P = 0.036). Faded terms fall between q = 0.051 and 0.058 and are shown for completeness. Asterisks mark genes also moved by A. muciniphila alone. **C.** The 32 restored genes with the largest Cd effects, separated by the direction of the Cd response and grouped by biological function. Each arrow runs from the Cd effect (red) to the effect under co-treatment (blue); ΔPSI is relative to vehicle, so an arrow ending at zero indicates complete restoration. Open circles show the effect of A. muciniphila alone on the same event, plotted irrespective of significance. The rMATS event class is beneath each gene symbol and % restoration is at the right, calculated as |ΔPSI (Cd+Akk vs. Cd)| / |ΔPSI (Cd vs. Vehicle)|, the size of the correction relative to the size of the Cd effect; values above 100% indicate that A. muciniphila moved the event past the vehicle baseline. Across these genes the median absolute effect of A. muciniphila alone was 0.104 ΔPSI, against 0.283 for Cd and 0.050 for the co-treated endpoint, and no gene reached significance for A. muciniphila alone at these events (FDR ≥ 0.05). Functional groups are descriptive; two correspond to the enriched terms in **B.**

**Table 2.** *A. muciniphila* reversed AS events ([Cd+Akk vs Cd] against [Cd vs. Vehicle]) and integration with RNA-Seq.

|  |  |  |  |  |  |  |
| --- | --- | --- | --- | --- | --- | --- |
| **AS type** | **\*Reversed events** | **Unique genes** | **<sup>†</sup>Overlapped DEGs** | **% DEGs** | **<sup>‡</sup>High-confidence events** | **High-confidence genes** |
| A3SS: alternative 3' splice site | 156 | 148 | 13 | **8.8%** | 22 | 21 |
| A5SS: alternative 5' splice site | 89 | 86 | 10 | **11.6%** | 12 | 12 |
| MXE: mutually exclusive exon | 80 | 76 | 4 | **5.3%** | 2 | 2 |
| RI: retained intron | 135 | 130 | 10 | **7.7%** | 7 | 7 |
| SE: skipped exon | **807** | **705** | **36** | **5.1%** | 56 | 51 |
| **Total** | **1,267** | **1,052** | **63** | **6.0%** | **99** | **92** |

<sup>\*</sup>Reversed events: splicing events changing significantly in opposite directions in Cd vs Vehicle and Cd+Akk vs Cd ( $p < 0.05$ in both comparisons).
<sup>†</sup>Overlapped DEGs: genes in the reversed set also differentially expressed by RNA-seq ( $p < 0.05$ ) in Cd vs Vehicle or Cd+Akk vs Cd. % DEGs is this count divided by unique genes.
<sup>‡</sup>High-confidence events and genes: reversed events additionally meeting $FDR < 0.05$ in both comparisons, a cadmium effect of at least 10% PSI, and no residual departure from the vehicle baseline (Cd+Akk vs. Vehicle, $FDR \geq 0.05$ ). High-confidence genes are the unique genes represented by those events.

To conduct an integrated triple-omics analysis and determine the extent to which post-transcriptional regulation was accompanied by changes in steady-state mRNA abundance, m⁶A epitranscriptomic and alternative-splicing (AS) datasets were integrated with RNA-seq results for each pairwise comparison (**Supplementary Table S2**). Cd vs Vehicle yielded 402 significant m⁶A events affecting 377 unique genes, of which 56 genes (14.9%) were also differentially expressed at the mRNA level. Akk vs Vehicle produced 1,075 significant m⁶A events across 979 genes, with 74 genes (7.6%) also showing significant mRNA-abundance changes. The largest number of m⁶A alterations was observed for Cd+Akk vs Cd, with 1,536 significant events affecting 1,425 genes; however, only 26 genes (1.8%) were also significant by RNA-seq. Thus, this analysis further confirmed that despite extensive m⁶A remodeling, particularly following Akk intervention in Cd-exposed animals, only a small fraction of m⁶A-regulated genes exhibited corresponding changes in total transcript abundance. **Table 3** further summarizes the triple-omics integration of m⁶A epitranscriptomics, alternative splicing (AS), and RNA-seq, identifying four unique hippocampal genes with coordinated reversal of both Cd-induced m⁶A modification and the same AS event following *A. muciniphila* intervention. Clavesin 2 (Clvs2) and leucine-rich repeat-containing 28 (Lrrc28) showed Cd-induced m⁶A hypermethylation that was reversed to hypomethylation by Akk, together with significant reversal of skipped-exon (SE) usage: ΔPSI changed from −0.039 to +0.051 for Clvs2 and from −0.070 to +0.055 for Lrrc28. In contrast, mucolipin 1 (Mcoln1) and transmembrane channel-like 4 (Tmc4) showed the opposite m⁶A pattern, with Cd-induced hypomethylation reversed to hypermethylation by Akk and were associated with retained-intron (RI) events. For Mcoln1, ΔPSI shifted from +0.016 to −0.028, whereas Tmc4 contained two independently reversed RI events, changing from −0.080 to +0.124 and from −0.073 to +0.064, respectively. Notably, none of these four genes was a significant RNA-seq DEG, indicating that the coordinated m⁶A and splicing reversals occurred without detectable changes in overall transcript abundance. Thus, these genes provide particularly strong examples of Akk-mediated restoration of Cd-disrupted post-transcriptional RNA regulation, while also suggesting that the relationship between m⁶A remodeling and splice-site usage is transcript– and event-specific rather than uniformly directional.

**Table 3.** Triple-omics integration: RNA-regulatory reversal by *A. muciniphila* without altered mRNA abundance.

|  |  |  |  |  |  |  |  |  |
| --- | --- | --- | --- | --- | --- | --- | --- | --- |
| Gene | m<sup>6</sup>A log<sub>2</sub>FC | Associated AS type | Cd vs Vehicle dPSI | Cd+Akk vs Cd dPSI | mRNA log<sub>2</sub>FC (Cd) | Hippocampal FPKM | Omics layer reversed | Nearest m<sup>6</sup>A site |
| *m<sup>6</sup>A hypermethylated under Cd, reversed by A. muciniphila* |  |  |  |  |  |  |  |  |
| Alkbh6 | +0.78 to -1.30* | SE | +0.038* | -0.004 | -0.04 | 36.2 | m<sup>6</sup>A only | ~1.3 kb (CDS) |
|  |  | SE | +0.061* | -0.008 |  |  |  |  |
| Clvs2 | +0.99 to -0.61* | SE | -0.039* | +0.051* | +0.06 | 23.7 | m<sup>6</sup>A + AS | overlap (5'UTR) |
| Lrrc28 | +0.91 to -0.60* | SE | -0.070* | +0.055* | -0.21 | 19.5 | m<sup>6</sup>A + AS | ~113 kb (CDS) |
| Pcsk4 | +1.68 to -1.40* | SE | -0.138*† | +0.116 | +0.37 | 2.5 | m<sup>6</sup>A only | overlap (CDS) |
| Spats1 | +0.67 to -0.89* | SE | -0.112* | +0.084 | +0.53 | 0.6 | m<sup>6</sup>A only | not reported |
| Tmc6 | +0.61 to -0.95* | A3SS | -0.365*† | -0.162 | +0.60* | 2.6 | m<sup>6</sup>A only | 1.5 kb (CDS) |
| Pgap2 | +0.99 to -1.01* | A5SS | -0.188* | +0.113 | +0.20 | 10.7 | m<sup>6</sup>A only | 25 kb (5'UTR) |
| *m<sup>6</sup>A hypomethylated under Cd, reversed by A. muciniphila* |  |  |  |  |  |  |  |  |
| Klhl13 | -1.33 to +0.71* | SE | -0.234* | +0.024 | -0.53* | 15.5 | m<sup>6</sup>A only | 47 kb (CDS) |
| Mcoln1 | -0.69 to +0.67* | RI (355 bp) | +0.016* | -0.028*† | +0.06 | 49.1 | m<sup>6</sup>A + AS | 2.9 kb (CDS) |
| Rhpn1 | -0.75 to +0.68* | RI (84 bp) | -0.106*† | +0.033 | -0.42 | 7.2 | m<sup>6</sup>A + AS | overlap (CDS) |
| Tmc4 | -0.67 to +0.92* | RI (93 bp) | -0.08*† | +0.124*† | -0.13 | 4.7 | m<sup>6</sup>A + AS | overlap (3'UTR) |
|  |  | RI (521 bp) | -0.073* | +0.064* |  |  |  |  |

\*Abbreviations and data interpretation. ΔPSI, difference in percent spliced in; FPKM, fragments per kilobase of transcript per million mapped reads. FPKM values represent vehicle-group mean baseline expression; roughly, 1 FPKM means low but detectable expression, and higher values indicate greater transcript abundance. These values are provided to show that the genes were detectable, not to indicate treatment-induced changes.
\*Statistical notation. \* $P < 0.05$ ; † FDR < 0.05; mRNA and m<sup>6</sup>A values.
mRNA abundance did not differ significantly for any gene shown ( $P = 0.36$ – $0.76$ ). m<sup>6</sup>A values are reported as log<sub>2</sub> fold changes in methylation, shown first for Cd versus vehicle and then for Cd+Akk versus Cd.
\*Gene info: Alkbh6, AlkB-family dioxygenase/metal-ion binding; Clvs2, lysosome/endosomal trafficking; Klhl13, Cul3 ubiquitin-ligase adaptor; Lrrc28, PML nuclear body protein; Mcoln1, lysosomal TRPML1 channel involved in iron transport/autophagy; Pcsk4, proprotein convertase/fertilization; Pgap2, GPI-anchor remodeling; Rhpn1, Rho GTPase effector/focal adhesion; Spats1, serine-rich nervous system/testis protein; Tmc4, chloride channel; Tmc6, mechanosensitive ER/Golgi ion channel.
\*Omics reversed lists the layers showing opposing changes under cadmium and *A. muciniphila* co-treatment. No gene showed a significant reversal of mRNA abundance ( $P \geq 0.09$ ), so mRNA does not appear in this column.
\*Nearest m<sup>6</sup>A site indicates the distance from the closest differentially methylated site to the regulated splicing event; “overlapping” means the site falls within the event bounds. Because genes contain 2–19 differentially methylated sites, proximity does not imply causality. Site coordinates were unavailable for Spats1.

To determine which regulatory layer carries the *A. muciniphila* response, we asked how often genes in two defined sets were altered at each layer (**Fig. 8**; gene-set membership and per-layer significance thresholds in **Supplementary Table S3**). Among genes encoding the RNA-regulatory machinery itself, Cd altered splicing in 31 of 72 alternative-splicing regulators (43.1%) and in 3 of 22 m⁶A regulators (13.6%), while the same gene sets showed m⁶A changes in at most 1 of 72 (1.4%) and abundance changes in at most 4 of 72 (5.6%). The pattern persisted under intervention, with splicing altered in 25 of 72 splicing regulators (34.7%) in the Cd+Akk vs. Cd comparison, against no m⁶A or abundance change in the m⁶A regulator set. Applying the same analysis to a combined set of 227 nonduplicated learning and memory and Alzheimer’s disease genes gave 28 genes altered by splicing (12.3%), 11 by m⁶A (4.8%) and 7 by abundance (3.1%). Alternative splicing therefore affected the largest fraction of genes in every comparison and in both gene sets, and the splicing machinery is itself a substrate of the program it controls.

**Figure 8.**
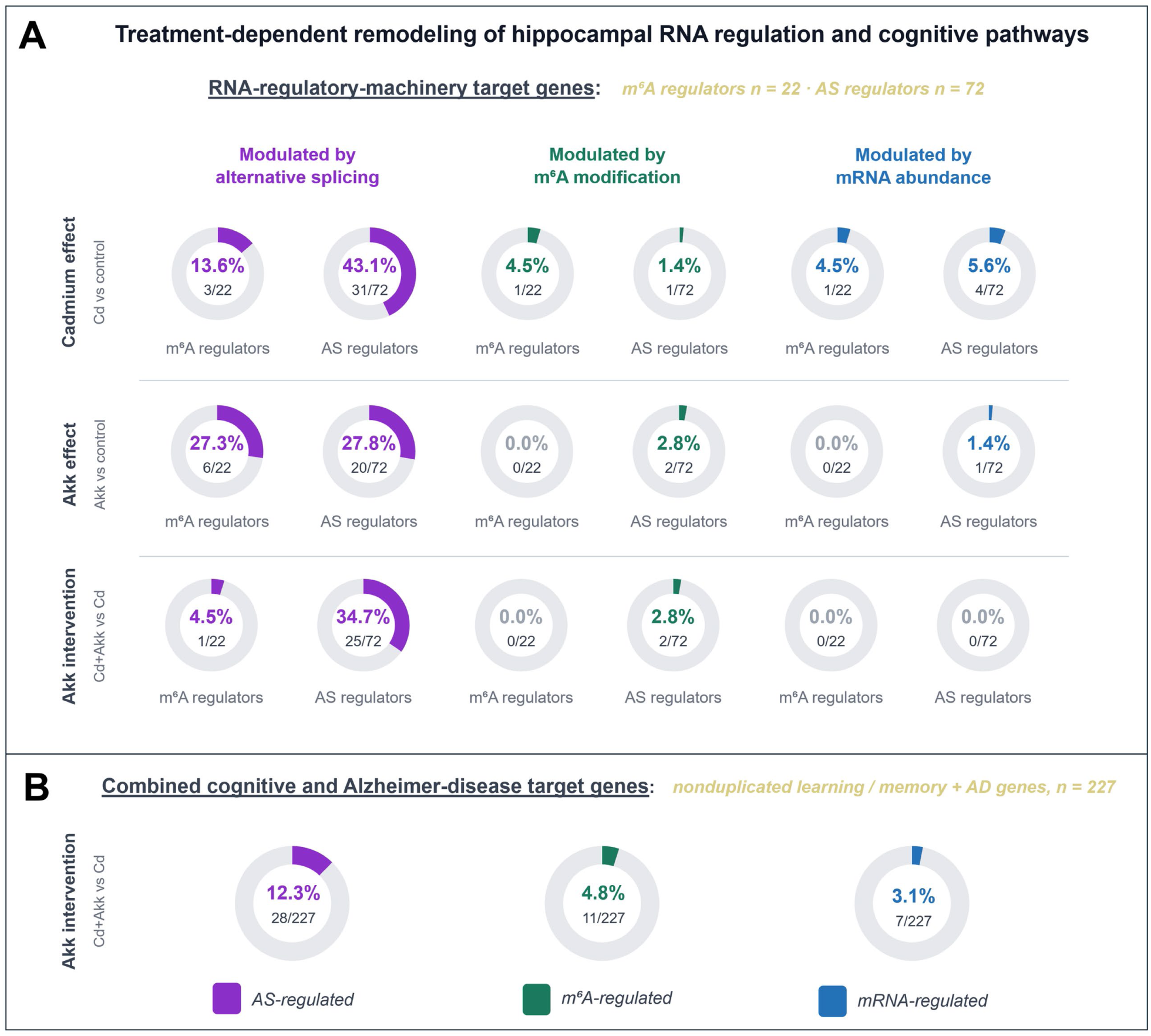
Treatment-dependent remodeling of hippocampal RNA regulation across splicing, m⁶A and abundance. Each ring shows the percentage of a defined gene set with a significant change in one regulatory layer: alternative splicing (purple), m⁶A modification (green) or mRNA abundance (blue); the fraction inside each ring is affected genes over the size of the set. A. Genes encoding RNA-regulatory machinery, comprising m⁶A regulators (n = 22, left ring of each pair) and alternative-splicing regulators (n = 72, right ring of each pair), evaluated for the cadmium effect (Cd vs. control), the A. muciniphila effect (Akk vs. control) and the A. muciniphila intervention (Cd+Akk vs. Cd). B. A combined set of nonduplicated learning and memory and Alzheimer’s disease genes (n = 227), evaluated for the A. muciniphila intervention only. Alternative splicing affected the largest fraction of genes in every comparison and in both gene sets, identifying it as the predominant regulatory layer engaged across both the RNA-regulatory machinery and cognitive and Alzheimer’s disease targets. Gene-set membership and the per-layer significance thresholds are given in Supplementary Table S3.

To determine what the restored splicing events do to the encoded proteins, each of the 99 high-confidence events was resolved to a transcript, exon and amino-acid interval and projected onto the AlphaFold model of its protein (Jumper et al., 2021), giving the structural triage shown in **Figure 9A**. Forty-nine of the 92 genes (53%) carry an event that introduces a premature termination codon and predicts nonsense-mediated decay, and therefore act on transcript fate rather than protein sequence (representative structures in **Supplementary Figure S3**); 34 genes (37%) carry a short in-frame edit within a loop or linker; and 9 genes (10%) carry a domain-scale in-frame edit (representative structures in **Supplementary Figure S4**). Eighty-three of the 92 restored genes thus alter transcript fate or make a short in-frame edit, and only nine carry a domain-scale edit, indicating that restoration operates predominantly by tuning transcript fate and flexible segments rather than by rebuilding folded domains. A restored ligand-receptor pair illustrates both mechanisms in one axis (**Fig. 9B**). In Plxnb3, Cd shifts the exon 2 acceptor and removes the Sema β-propeller that holds B-type plexins inactive (amino acids 5-314, in-frame, FDR 4.8 × 10⁻³); because this domain is autoinhibitory, its loss is predicted to yield a ligand-independent receptor rather than a non-functional one (Takahashi and Strittmatter, 2001). In Sema4a, Cd shifts the exon 6 acceptor and removes a 47-nucleotide segment that introduces a premature termination codon, so that less of the transcript is degraded and semaphorin output rises (FDR 4.1 × 10⁻²). *A. muciniphila* restored both events, by 97% and 99% respectively, and the two proteins act across a single cell-cell interface, with Sema4A presented by activated microglia and lymphocytes (Leitner et al., 2015; Koda et al., 2020) and Plexin-B3 selectively expressed in white matter and postnatal oligodendrocytes (Worzfeld et al., 2004). Sema4A binds all three B-type plexins (Yukawa et al., 2010) and, in the developing hippocampus, promotes inhibitory and excitatory synapse formation through Plexin-B1 and Plexin-B2 (McDermott et al., 2018), so increased Sema4A output could also act on hippocampal neurons directly. Four additional in-frame switches mediated by *A. muciniphila* are shown for Dctn1, Efna5, Ncoa6 and Phactr4) in **Figure 9C**. The affected sequences map to regions associated with axonal transport, ephrin receptor binding, transcriptional activation, and PP1/actin regulation, respectively. These annotations suggest possible functional effects but have not been validated at the protein level. The structure-function call for all significant, restored splicing event is listed in **Supplementary Table S1**.

**Figure 9.**
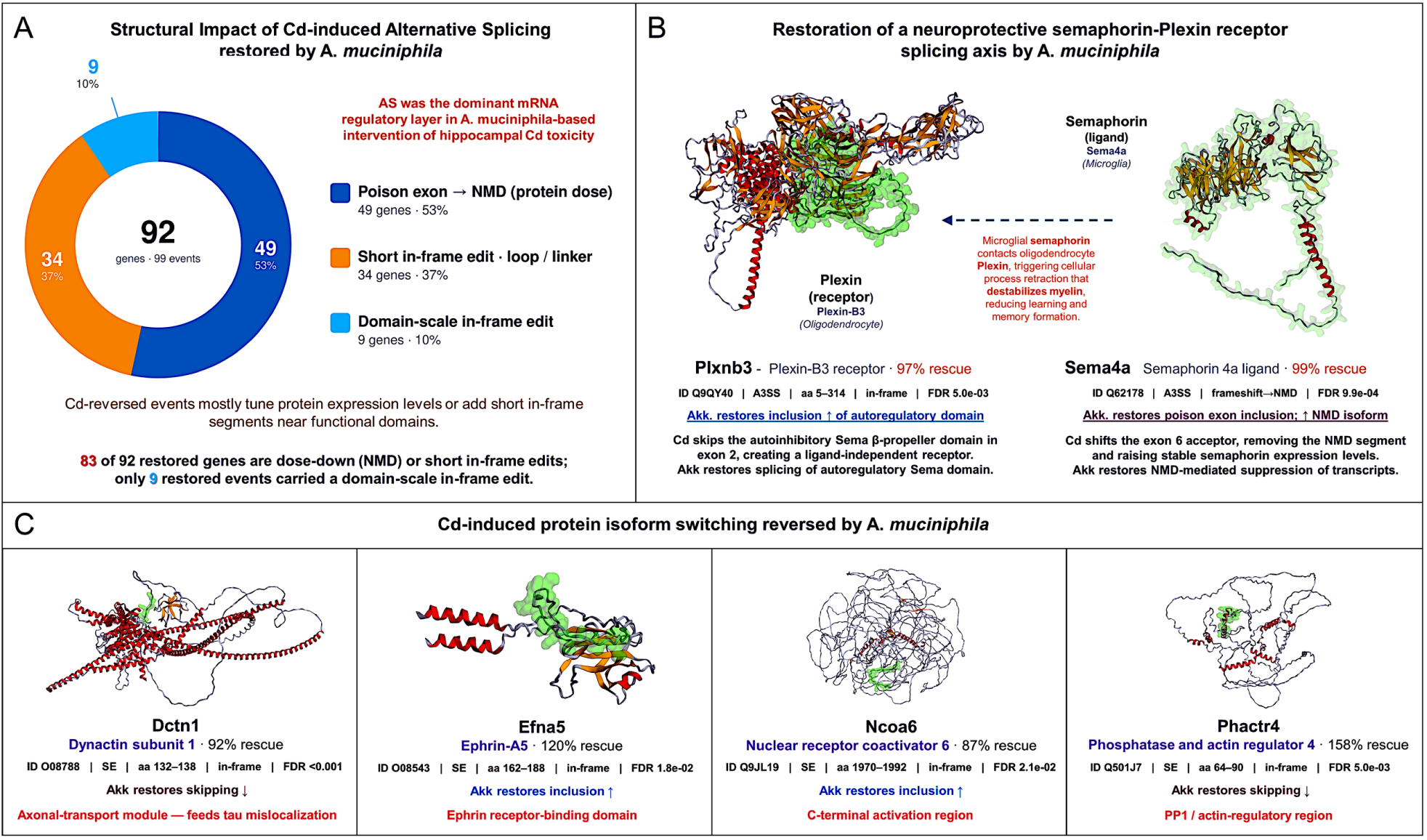
Predicted structural consequences of Cd-disrupted splicing restored by A. muciniphila. **A.** The 99 high-confidence restored events in 92 genes were classified by their predicted effect on transcript fate or protein structure (Supplementary Table S1). Events in 49 genes (53%) are predicted to introduce a premature stop and trigger nonsense-mediated decay; 34 (37%) produce short in-frame edits in loops or linkers; and nine (10%) produce domain-scale in-frame edits. Thus, most restored genes are predicted to change transcript fate or a short protein segment rather than an entire folded domain. **B.** Sema4a and Plxnb3 illustrate two splice changes in a potential cell-contact signaling axis: Sema4A is a ligand for B-type plexins, including Plexin-B3, which is expressed in oligodendrocytes (Worzfeld et al., 2004; Yukawa et al., 2010). Cd favors a Plxnb3 isoform lacking its Sema β-propeller and exclusion of a 47-nt Sema4a segment predicted to generate a premature stop when retained. A. muciniphila reverses both changes, reducing their absolute deviation from vehicle by 97% and 99%, respectively. The proposed effects on receptor activity and Sema4A output remain untested. **C.** Additional restored in-frame segments map to an axonal-transport region in DCTN1, a 27-amino-acid EFNA5 spacer adjacent to the Eph receptor-binding domain, the NCOA6 C-terminal activation region, and a PP1/actin-regulatory region in PHACTR4. Green highlights the affected segments on AlphaFold models. Percent restoration in C and Supplementary Table S1 measures the Akk-induced correction relative to the Cd effect; values above 100% indicate overshoot of vehicle baseline.

Taken together, building on from our previous publication that *A. muciniphila* prevented Cd-induced memory deficit accompanied with microbiome and metabolome reprogramming (Wang et al., 2026a), as shown in **Fig. 8**, the present study used the same mouse cohort and demonstrated that Cd exposure disrupts hippocampal RNA regulation across multiple molecular features, producing transcriptional alterations in neuronal, neurovascular, inflammatory, metabolic, and stress-response programs together with substantially broader changes in m⁶A epitranscriptomic regulation and alternative splicing. In contrast, *A. muciniphila* intervention opposed key Cd-associated molecular alterations, including directional reversal of 12 Cd-responsive transcripts, 53 unique genes with reversed m⁶A states, 45 of which reverse at the level of the same transcript in both comparisons, and 1,052 genes with reversal of the same Cd-induced alternative-splicing events, with functional signatures converging on neuronal plasticity and excitability, synaptic transmission, neuroimmune regulation, cellular stress/genome maintenance, and metabolic homeostasis. Importantly, integration of the three omics layers revealed that only a small fraction of m⁶A-or AS-regulated genes exhibited corresponding changes in steady-state mRNA abundance, and several genes showed coordinated reversal of m⁶A and splice-isoform regulation in the complete absence of differential total mRNA expression. Together, these findings suggest that Cd neurotoxicity extends beyond transcriptional dysregulation to disruption of hippocampal post-transcriptional RNA architecture, whereas the protective effect of *A. muciniphila* is associated predominantly with restoration of m⁶A regulation and splice-isoform selection rather than broad normalization of transcript abundance. These molecular findings provide a potential mechanistic framework for the previously observed protection against Cd-induced memory deficits in the same mouse cohort, while highlighting the need for future proteomic, translational, and isoform-specific functional studies to determine how these rescued RNA regulatory states translate into protein expression, neuronal function, and cognitive outcomes.

## DISCUSSION

The present study advances knowledge of the gut-brain axis and of remediation against Cd neurotoxicity by showing that *A. muciniphila* supplementation reverses Cd-responsive genes at the transcriptome, m⁶A epitranscriptome, and alternative-splicing levels. It also points to a previously under-appreciated gut-brain mechanism in which the neuroprotection reported earlier (Wang et al., 2026a) is linked predominantly to restoration of hippocampal post-transcriptional RNA architecture, particularly splice-isoform selection and m⁶A regulation, rather than to altered overall mRNA abundance.

These molecular findings build on our recent report using the same mouse cohort, in which oral *A. muciniphila* supplementation prevented Cd-induced memory impairment without reducing brain Cd concentrations (Wang et al., 2026a), indicating that the effect is unlikely to reflect altered Cd toxicokinetics. The present results indicate that reprogramming of the hippocampal RNA regulatory network may contribute to that protection, identifying the gut-brain axis as a target in neurotoxicity. This interpretation is supported by our earlier demonstration that environmentally relevant Cd exposure induces gut dysbiosis before the onset of cognitive impairment (Wang et al., 2025), and by reports that *A. muciniphila* improves cognition and reduces cerebral amyloid pathology in APP/PS1 mice (Ou et al., 2020), improves cognition in aged mice by suppressing IL-6-associated inflammation (Zhu et al., 2023), and modulates hippocampal GABA/glutamate balance (Olson et al., 2018). Consistent with these gut-brain effects, RNA-seq identified 12 Cd-responsive hippocampal transcripts whose directional changes were opposed by *A. muciniphila*, converging at the pathway level on neuropeptide signaling, neuronal excitability, mitochondrial function, lipid homeostasis, proteostasis, and redox defense, processes that collectively govern hippocampal resilience (Lee et al., 2018; Smith et al., 2020; Guo et al., 2021; Tseng et al., 2023). The predicted enrichment of Nrf2-centered cytoprotective signaling among the reversed transcripts is notable, because Nrf2 is a central regulator of antioxidant and detoxification programs (Yamamoto et al., 2018) and restrains Cd-induced microglial overactivation, synaptic engulfment, and learning-memory impairment (Li et al., 2025). Rather than preventing Cd accumulation in the brain, *A. muciniphila* thus appears to reprogram hippocampal neuronal, inflammatory, metabolic, and antioxidant-response networks, linking the microbiome intervention to the behavioral neuroprotection observed previously.

A particularly striking finding was the magnitude of *A. muciniphila*-mediated restoration of Cd-disrupted alternative splicing: 1,267 Cd-induced AS events in 1,052 unique genes were reversed by Akk, with skipped-exon events the largest class (Table 2). Only 63 of these genes (6.0%) also showed significant changes in total mRNA abundance, indicating that isoform restoration occurred largely independently of gene-level transcription. Applying FDR correction across both comparisons together with a requirement for return to the vehicle baseline narrows this to 99 events in 92 genes, the high-confidence core of the splicing response (Figure 7, Supplementary Table S1). This core is enriched for regulators of mRNA 3′-UTR AU-rich element binding (*Arid5a*, *Cpeb1*, *Cpeb2*, *Exosc8*; adjusted P = 2.3 × 10⁻⁵) and for calcium-induced calcium release (*Ryr1*, *Ryr3*; adjusted P = 0.017), placing the restored events on machinery governing local translation and calcium-dependent plasticity. The same genes recurred across four of the nine enriched terms, indicating a coherent post-transcriptional module rather than independent signals. Because these proteins act on 3′ untranslated regions rather than on transcription, their restoration would act downstream of transcript synthesis. This matters because alternative splicing is an exceptionally prominent regulatory mechanism in the nervous system, controlling neuronal differentiation, synapse specification, and network function (Traunmuller et al., 2016; Furlanis and Scheiffele, 2018; Traunmuller et al., 2023); isoform choice alone can set synaptic strength, as for GluN1 in hippocampal long-term potentiation and spatial memory (Sengar et al., 2019). Consistent with this, our 63 RNA-seq-overlapping AS-reversal genes were enriched for excitatory postsynaptic potential, neurotransmitter secretion, synaptic transmission, neuronal projection organization, and Wnt signaling. Splicing regulation can be dissociated from transcript abundance in the human brain (Tollervey et al., 2011), and normalization of splicing programs has been associated with neuroprotection in AD-related tauopathy (Apicco et al., 2019). Notably, *Clvs2*, *Lrrc28*, *Mcoln1*, and *Tmc4* showed coordinated reversal of both Cd-induced m⁶A alterations and the corresponding AS events, despite no significant change in total mRNA abundance. These associations do not establish that m⁶A caused the splicing changes, but such coupling is mechanistically plausible: the nuclear m⁶A reader YTHDC1 regulates exon selection through splicing factors (Xiao et al., 2016), and neuronal METTL3/m⁶A depletion produces aberrant splicing, including exon exclusion in synapse-associated transcripts (Wang CX et al., 2018).

A second major finding is that m⁶A remodeling may serve as an important epitranscriptomic component of *A. muciniphila*-mediated remediation of Cd neurotoxicity. Akk intervention reversed Cd-induced m⁶A changes in 53 unique hippocampal genes, spanning immune and neurovascular signaling, oxidative stress and genome maintenance, synaptic plasticity and FGF-MAPK signaling, lysosome-autophagy regulation, and nucleotide metabolism. Representative genes support these annotations: the presynaptic organizer *Fgf22* (Terauchi et al., 2016), the neural-stem-cell regulator *Tsku* (Ahmad et al., 2020), *Spp1*, which mediates microglial synaptic engulfment in AD mouse hippocampus (De Schepper et al., 2023), the lysosomal ROS sensor *Mcoln1* (Zhang et al., 2016), and the redox-coupled differentiation factor *Prdx4* (Yan et al., 2015). Importantly, this extensive m⁶A response was largely uncoupled from steady-state transcript abundance: in the Cd+Akk vs Cd comparison, only 26 of 1,425 genes with significant m⁶A changes (1.8%) were also differentially expressed. This disconnect is mechanistically plausible, because m⁶A determines RNA fate rather than simply transcript quantity, directing turnover, translation efficiency, splicing, nuclear export, and the localized synaptic epitranscriptome through distinct readers (Wang et al., 2014; Wang et al., 2015; Xiao et al., 2016; Roundtree et al., 2017; Merkurjev et al., 2018). Limited correspondence with mRNA abundance is therefore not evidence of a lack of functional consequence; it raises the possibility that *A. muciniphila* restores the post-transcriptional fate and translational competence of Cd-disrupted transcripts without changing their abundance, a hypothesis requiring proteomics and transcript-specific functional studies.

The structural impact of Cd-induced alternative splicing discussed in **Figure 9** constrains how this restoration is likely to act. Because 83 of the 92 high-confidence genes either alter transcript fate through nonsense-mediated decay or make a short in-frame edit in a loop or linker, and only nine carry a domain-scale edit, the splicing response is better described as retuning to the basal amount and nature of existing proteins, rather than as repairing damaged or compromised domains. Framing the restored set this way also identifies which events are tractable for splice-switching antisense oligonucleotides, the strategy we previously applied to cytochrome P450 isoform selection (Annalora et al., 2020; Lidberg et al., 2021). That distinction matters, because changes in transcript fate and disordered-segment edits are least likely to be visible in gene-level expression data, consistent with the limited overlap observed. The semaphorin-plexin pair is informative for a second reason: both changes increase signaling through the same axis, one by removing the autoinhibitory domain of the receptor and the other by raising ligand output, and both are reversed by *A. muciniphila*. Because Sema4A is cytotoxic to oligodendrocytes and drives Th17-mediated neuroinflammation (Leitner et al., 2015; Koda et al., 2020), while adult oligodendrogenesis and new myelin formation are required for hippocampal memory consolidation (Steadman et al., 2020; Pan et al., 2020; Xin and Chan, 2020), excess signaling through this axis offers a testable route from a splicing change to the memory phenotype. We emphasize that these are structural predictions from transcript-level measurements only; neither protein abundance, receptor localization, nor myelin integrity was measured here, and the consequences of removing the Plexin-B3 Sema domain, including whether the truncated receptor reaches the membrane, remain to be established.

Several limitations apply to our integrated triple-Omics study overall. First, this study defines RNA-level regulatory signatures rather than their functional consequences. Neither quantitative proteomics nor direct measurements of translation were performed, so it remains unknown whether reversal of m⁶A states or AS events translates into changes in protein abundance, isoform composition, localization, or activity; proteomics and ribosome profiling will be needed. Similarly, the coordinated reversal of m⁶A and splicing is associative; studies manipulating m⁶A writers, erasers, or readers will be required to establish causality. Second, only male mice were included; young male mice were previously more vulnerable than females to Cd-associated cognitive impairment in an ApoE4 model (Zhang et al., 2020). Because sex influences Cd toxicodynamics, microbiome composition, and neuroimmune responses, future studies should evaluate sex-dependent responses. Third, species differences in microbiome composition, xenobiotic metabolism, brain physiology, and m⁶A regulation limit translation to humans; validation in human neuronal or organoid models will be important. Finally, bulk tissue precludes cell-type resolution, and the exploratory use of nominal p < 0.05 increases sensitivity at the cost of possible false positives.

Despite these limitations, the present study demonstrates that the neuroprotective effects of *A. muciniphila* against Cd exposure are associated with coordinated remodeling of the hippocampal transcriptome, m⁶A epitranscriptome, and alternative-splicing landscape. By integrating three complementary layers of RNA regulation in the same model in which *A. muciniphila* prevented Cd-induced memory impairment (Wang et al., 2026a), this work shows that a substantial component of the response occurs post-transcriptionally, often without corresponding changes in total mRNA abundance. Structurally, most restored events act on transcript fate or edit short, flexible protein segments rather than rebuilding folded domains, and the semaphorin-plexin axis illustrates how such a change could reach the memory phenotype. The convergence of these alterations on neuronal signaling, synaptic function, redox homeostasis, and RNA processing provides a framework for how microbial intervention may influence brain resilience, and a foundation for targeting gut-brain epitranscriptomic regulation to mitigate environmentally associated neurodegenerative risk.

## DATA AVAILABILITY

Raw and processed data generated in this study (RNA-seq, m⁶A epitranscriptomic microarray, and rMATS alternative-splicing output) will be deposited in the NCBI Gene Expression Omnibus (GEO) and publicly released upon publication. All additional data supporting the findings of this study are available in the Supplementary Material or from the corresponding authors upon reasonable request.

## ACKNOWLEDGEMENTS

The authors thank the Oregon State University Center for Quantitative Life Sciences (CQLS) for computational infrastructure and sequencing support, and members of the Cui Laboratory for proofreading the manuscript.

## AUTHOR CONTRIBUTIONS

Z.K., conceptualization, data analysis, investigation, visualization, manuscript preparation;

J.L., data curation, data analysis, software;

A.J.A., data analysis, methodology, software, visualization, manuscript preparation;

H.W., conceptualization, investigation, conducting the laboratory experiments, review and editing of the manuscript;

J.Y.C., conceptualization, investigation, conducting the laboratory experiments, funding acquisition, project administration, supervision, manuscript preparation, review and editing.

