## Supplementary Material for "Triple-Omics Analysis Unveiled the Neuroprotective Mechanisms of *Akkermansia muciniphila* in an Environmental Exposure-Linked Mouse Model of Alzheimer’s Disease"

### Contents

Supplementary Figure S1. Transcript abundance in genes whose splicing was reversed by *A. muciniphila*

Supplementary Figure S2. Splice-isoform changes occur largely independently of transcript abundance

Supplementary Figure S3. Restored poison-exon events predicted to act through nonsense-mediated decay

Supplementary Figure S4. Structural classes of A. muciniphila-restored splicing events

Supplementary Table S1. High-confidence restoration of Cd-disrupted alternative splicing by *A. muciniphila*

Supplementary Table S2. mRNA expression-anchored data integration with $m^6A$ and AS

Supplementary Table S3. Gene-set membership and per-layer significance thresholds (Figure 8)

**RNA-seq change in the 63 genes with A. muciniphila-reversed splicing**

| Event type | Gene | Cd vs vehicle | p | Cd + Akk vs Cd | p |
|---|---|---|---|---|---|
| A3SS – Alternative 3′ splice site (13 genes) | *Ankrd13c* | **−0.21** | p = 0.046 | +0.09 | p = 0.423 |
| | *Arntl2* | **+1.42** | p < 0.001 | −0.29 | p = 0.182 |
| | *Camk2b* | **+0.25** | p = 0.042 | +0.01 | p = 0.939 |
| | *Camk2d* | **−0.35** | p = 0.019 | **+0.49** | p = 0.003 |
| | *Cdh18* | **−0.44** | p = 0.009 | +0.02 | p = 0.924 |
| | *Kcnc3* | **+0.30** | p = 0.005 | −0.16 | p = 0.145 |
| | *Neurl4* | **+0.27** | p = 0.031 | −0.09 | p = 0.515 |
| | *Nfix* | **+0.29** | p = 0.007 | −0.13 | p = 0.256 |
| | *Nudcd2* | **+0.57** | p = 0.026 | −0.32 | p = 0.180 |
| | *Osbpl5* | **−0.40** | p = 0.034 | +0.07 | p = 0.714 |
| | *Pkp4* | −0.03 | p = 0.781 | **−0.27** | p = 0.027 |
| | *Tmem108* | **+0.32** | p = 0.017 | 0.00 | p = 0.980 |
| | *Zfp414* | **+0.30** | p = 0.044 | −0.15 | p = 0.312 |
| A5SS – Alternative 5′ splice site (7 genes) | *Atp13a4* | **+0.52** | p = 0.020 | −0.42 | p = 0.065 |
| | *Cep89* | **+0.46** | p = 0.047 | −0.10 | p = 0.678 |
| | *Fgfr1* | **+0.25** | p = 0.023 | −0.02 | p = 0.826 |
| | *Lamp5* | **−0.25** | p = 0.033 | +0.07 | p = 0.551 |
| | *Ngb* | **−0.59** | p = 0.014 | **+0.58** | p = 0.019 |
| | *Tcf7l2* | **−1.35** | p < 0.001 | −0.40 | p = 0.111 |
| | *Wnk1* | **−0.40** | p = 0.005 | +0.26 | p = 0.075 |
| MXE (3 genes) | *Ccdc162* | +0.75 | p = 0.089 | **−1.03** | p = 0.021 |
| | *P4ha1* | **−0.35** | p = 0.001 | +0.14 | p = 0.239 |
| | *Sulf1* | −0.15 | p = 0.400 | **−0.58** | p = 0.002 |
| RI – Retained intron (10 genes) | *Adamts10* | **+0.30** | p = 0.033 | −0.05 | p = 0.751 |
| | *Amdhd2* | **+0.46** | p = 0.009 | −0.12 | p = 0.494 |
| | *Baiap3* | **−1.00** | p < 0.001 | +0.59 | p = 0.072 |
| | *Ergic3* | **−0.31** | p = 0.021 | +0.15 | p = 0.225 |
| | *Mtmr2* | **−0.31** | p = 0.008 | −0.02 | p = 0.851 |
| | *Pcolce* | **+0.42** | p = 0.031 | **−0.46** | p = 0.023 |
| | *Psmb9* | **+1.22** | p = 0.028 | −1.01 | p = 0.054 |
| | *Ptpn5* | **−0.24** | p = 0.011 | +0.10 | p = 0.286 |
| | *Trp53i11* | −0.01 | p = 0.955 | **+0.26** | p = 0.013 |
| | *Zcchc9* | **−0.55** | p = 0.022 | +0.16 | p = 0.489 |

| Event type | Gene | Cd vs vehicle | p | Cd + Akk vs Cd | p |
|---|---|---|---|---|---|
| SE – Skipped exon (30 genes) | *Adamtsl2* | **+0.67** | p = 0.023 | −0.10 | p = 0.708 |
| | *Adgrl2* | **−0.25** | p = 0.037 | +0.17 | p = 0.183 |
| | *Ap4e1* | −0.20 | p = 0.279 | **+0.42** | p = 0.033 |
| | *Cxxc4* | **−0.29** | p = 0.016 | +0.01 | p = 0.930 |
| | *Dbndd2* | **−0.29** | p = 0.036 | +0.18 | p = 0.285 |
| | *Epn1* | **+0.26** | p = 0.009 | −0.04 | p = 0.680 |
| | *Fhod1* | **−0.84** | p = 0.030 | +0.04 | p = 0.921 |
| | *Grik4* | **+0.34** | p = 0.003 | −0.08 | p = 0.473 |
| | *Ivns1abp* | **−0.26** | p = 0.011 | 0.00 | p = 0.998 |
| | *Lrrk2* | **−0.50** | p = 0.012 | +0.04 | p = 0.857 |
| | *Meis3* | **+0.29** | p = 0.047 | −0.23 | p = 0.119 |
| | *Mthfs* | −1.16 | p = 0.265 | **−1.60** | p < 0.001 |
| | *Myo5b* | **+0.30** | p = 0.038 | −0.06 | p = 0.690 |
| | *Nbeal2* | +0.06 | p = 0.797 | **+0.58** | p = 0.013 |
| | *Nkain4* | **+0.37** | p = 0.009 | −0.26 | p = 0.081 |
| | *Ntng1* | **−0.32** | p = 0.017 | +0.01 | p = 0.945 |
| | *Ntrk3* | **+0.55** | p = 0.002 | −0.24 | p = 0.118 |
| | *Pcbp4* | **+0.22** | p = 0.037 | +0.02 | p = 0.859 |
| | *Phldb1* | +0.12 | p = 0.399 | **−0.29** | p = 0.039 |
| | *Plch2* | **+0.27** | p = 0.009 | +0.03 | p = 0.750 |
| | *R3hdm2* | **−0.26** | p = 0.029 | +0.14 | p = 0.239 |
| | *Rgs17* | **−0.24** | p = 0.017 | +0.05 | p = 0.614 |
| | *Rnf13* | **−0.21** | p = 0.039 | −0.05 | p = 0.611 |
| | *Slc38a2* | **−0.27** | p = 0.005 | +0.05 | p = 0.611 |
| | *Sp100* | **+1.18** | p = 0.034 | −0.44 | p = 0.353 |
| | *Syt9* | **−0.51** | p = 0.007 | −0.19 | p = 0.350 |
| | *Taz* | **+0.35** | p = 0.011 | −0.12 | p = 0.388 |
| | *Tbl1xr1* | −0.32 | p = 0.070 | **−0.40** | p = 0.019 |
| | *Wiz* | **+0.28** | p = 0.038 | −0.02 | p = 0.862 |
| | *Zeb1* | −0.08 | p = 0.568 | **+2.16** | p < 0.001 |

−2 −1 0 +1 +2

**RNA-seq log2 fold change**

Bold values: p < 0.05 (Cuffdiff, unadjusted); grey values: not significant.

Genes in red are significant and opposite in direction in both comparisons.

**Supplementary Figure S1. Transcript abundance in genes whose splicing was reversed by *A. muciniphila*.** The 63 genes with a reversed splicing event that were also differentially expressed at nominal $p < 0.05$ in Cd vs. Vehicle or Cd+Akk vs. Cd, grouped by the representative reversed event type (A3SS, alternative 3′ splice site; A5SS, alternative 5′ splice site; MXE, mutually exclusive exons; RI, retained intron; SE, skipped exon). Genes with more than one reversed event are assigned to the most significant. Color shows the log2 fold change in transcript abundance for each comparison, with the Cuffdiff p value beside it; bold values are $p < 0.05$. Three genes, *Camk2d*, *Ngb* and *Pcolce* (red), are significant and opposite in direction in both comparisons

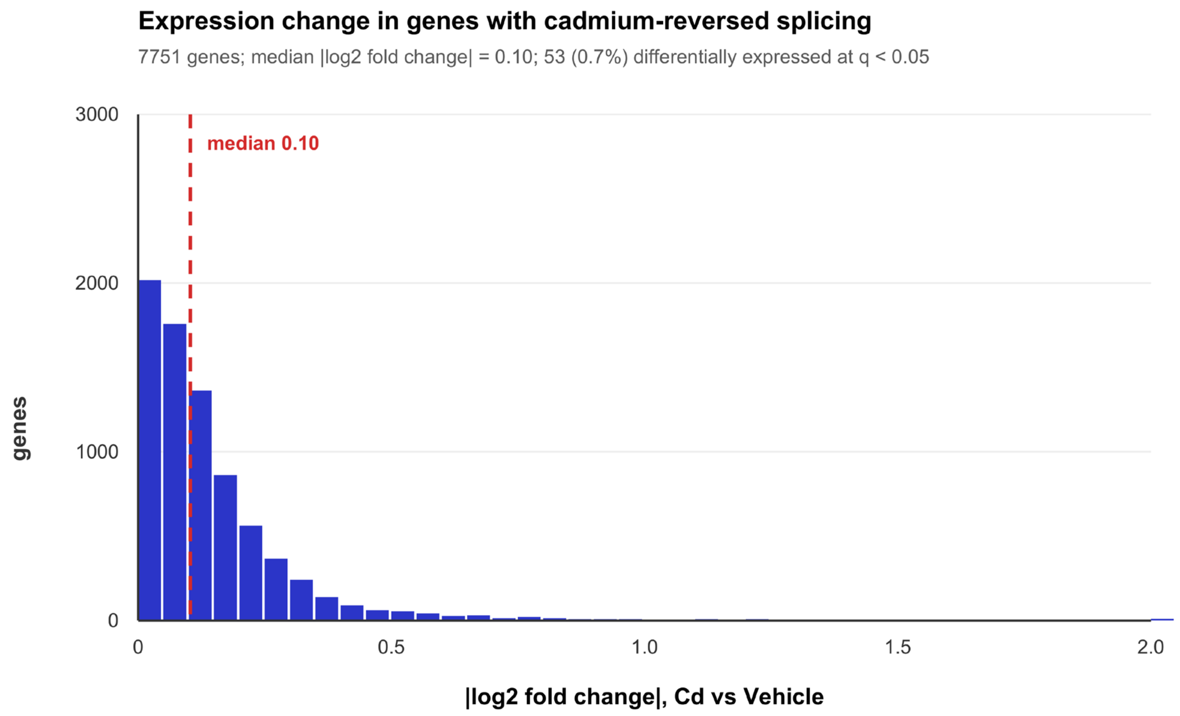


**Supplementary Figure S2. Splice-isoform changes occur largely independently of transcript abundance.** Distribution of |log2 fold change| in transcript abundance under Cd (Cd vs. Vehicle) for all 7,751 genes with usable RNA-seq data that carry at least one splicing event whose ΔPSI changed sign between Cd vs. Vehicle and Cd+Akk vs. Cd. No significance threshold is applied to the splicing events in this panel; the significance-filtered reversal set is described in the main text. The median is 0.10 (dashed line), and 53 genes (0.7%) were differentially expressed at $q < 0.05$.

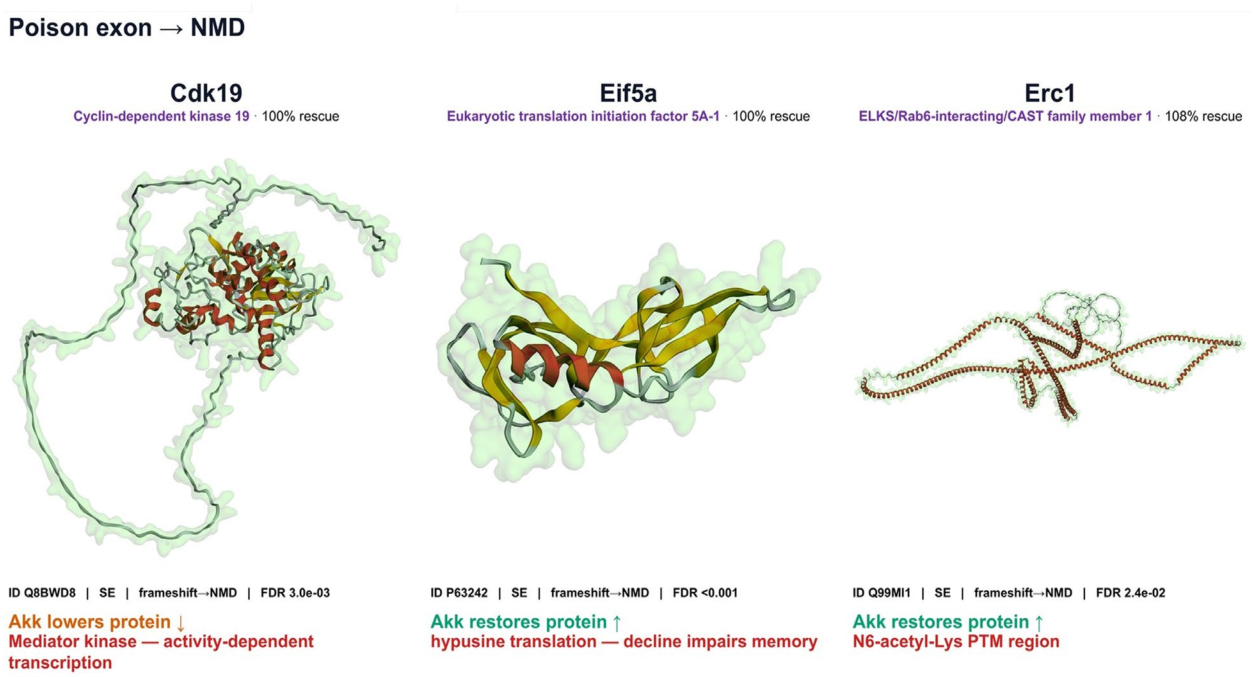


**Supplementary Figure S3. Restored poison-exon events predicted to act through nonsense-mediated decay**. Three of the 49 high-confidence restored genes whose reversed segment introduces a premature termination codon, shown on their AlphaFold models with the affected segment highlighted. Cdk19 (cyclin-dependent kinase 19) is a Mediator-complex kinase supporting activity-dependent transcription; Eif5a (eukaryotic translation initiation factor 5A-1) acts in hypusine-dependent translation, whose decline impairs memory; and Erc1 (ELKS/Rab6-interacting/CAST family member 1) is an active-zone scaffold for synaptic transmission. Percent restoration is |ΔPSI(Cd+Akk vs. Cd)| / |ΔPSI(Cd vs. Vehicle)|: Cdk19 and Eif5a are fully restored (100%) and Erc1 carries the event slightly past the vehicle baseline (108%). Because these events alter protein dose rather than protein sequence, *A. muciniphila* acts here on the amount of protein produced rather than on its folded structure.

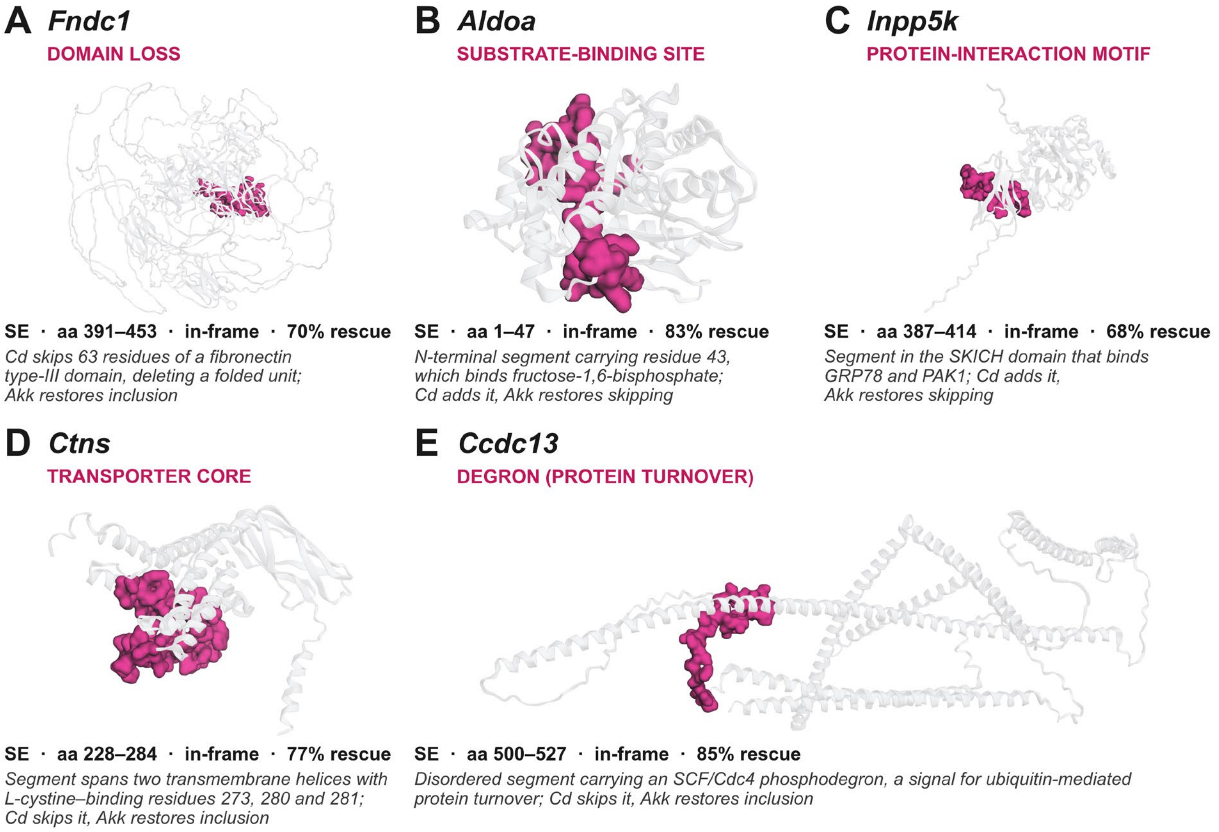


**Supplementary Figure S4. Structural classes of *A. muciniphila*-restored splicing events.** Representative in-frame events from the 99 high-confidence restored events in 92 genes (Supplementary Table S1), shown on AlphaFold models of the encoded proteins (translucent cartoon) with the Cd-switched segment as a magenta surface. Each panel gives the event type, amino-acid interval, reading frame and percent rescue by *A. muciniphila*.

**Supplementary Table S1. High-confidence restoration of Cd-disrupted alternative splicing by *A. muciniphila*.**

Ninety-nine splicing events in 92 genes meeting all four criteria: FDR < 0.05 in both Cd vs. Vehicle and Cd+Akk vs. Cd, opposite direction between those comparisons, |ΔPSI| ≥ 0.10 under Cd, and no residual departure from the vehicle baseline (Cd+Akk vs. Vehicle, FDR ≥ 0.05). Rows are grouped by the direction of the Cd effect, then alphabetically. % restoration is |ΔPSI(Cd+Akk vs. Cd)| / |ΔPSI(Cd vs. Vehicle)|, the size of the correction relative to the size of the Cd effect: 100% means the correction exactly offsets the Cd effect, below 100% means it falls short, and above 100% means A. muciniphila carried the event past the vehicle baseline in the opposite direction (overshoot). 35 of the 92 events overshoot on this measure. No event was corrected in the same direction as the Cd effect.

| Gene | Event type | Reversed segment (aa) | Structure-function call | Baseline PSI | Cd vs Veh ΔPSI | Cd FDR | Cd+Akk vs Cd ΔPSI | Cd+Akk vs Cd FDR | Cd+Akk vs VehΔPSI | % restoration | Functional group | Figure 7 group | Akk-alone effect | Cd-altered events (of total in gene) |
|---|---|---|---|---|---|---|---|---|---|---|---|---|---|---|
| | | | | | | | ***Cd increased inclusion (n = 42)*** | | | | | | | |
| ***Adamts6*** | SE | 611-644 | Poison exon → NMD | 0.471 | +0.505 | 0.022 | -0.359 | 0.001 | +0.146 | 71% | Metabolism, transport and cell structure | | same direction as Cd | 1 of 3 |
| ***Aldoa*** | SE | 1-47 | In-frame · Functional-element (F-1,6-BP site) | 0.110 | +0.155 | <0.001 | -0.129 | 0.010 | +0.026 | 83% | Metabolism, transport and cell structure | | none | 1 of 7 |
| ***Ankrd17*** | A3SS | — | In-frame · Loop/Disordered | 0.840 | +0.128 | 0.006 | -0.301 | <0.001 | -0.173 | 235% | Transcription, chromatin and RNA processing | | none | 1 of 5 |
| ***Arid5a*** | A3SS | — | In-frame · Loop/Disordered | 0.689 | +0.184 | 0.004 | -0.118 | 0.007 | +0.066 | 64% | Transcription, chromatin and RNA processing | RNA processing and translation | none | 2 of 4 |
| ***Atad5*** | A5SS | — | In-frame · Loop/Disordered | 0.623 | +0.321 | 0.018 | -0.420 | 0.010 | -0.099 | 131% | Proteostasis, DNA damage and stress response | Ubiquitin, proteostasis and repair | none | 1 of 3 |
| ***Atp13a4*** | A5SS | — | Poison exon → NMD | 0.835 | +0.133 | 0.020 | -0.153 | 0.010 | -0.020 | 115% | Metabolism, transport and cell structure | | none | 1 of 10 |
| ***Cbll1*** | A3SS | — | In-frame · Loop/Disordered | 0.807 | +0.137 | 0.016 | -0.126 | 0.012 | +0.010 | 92% | Transcription, chromatin and RNA processing | RNA processing and translation | none | 1 of 4 |
| ***Ccser2*** | A5SS | — | In-frame · Loop/Disordered | 0.057 | +0.173 | 0.001 | -0.182 | <0.001 | -0.009 | 105% | Metabolism, transport and cell structure | | none | 1 of 7 |
| ***Cdh18*** | A3SS | — | In-frame · Loop/Disordered | 0.422 | +0.578 | <0.001 | -0.627 | 0.030 | -0.049 | 108% | Synaptic signaling / neuronal plasticity | Calcium and cytoskeletal signaling | none | 2 of 7 |
| ***Cdk19*** | SE | 153-171 | Poison exon → NMD | 0.832 | +0.129 | 0.003 | -0.129 | 0.001 | +0.000 | 100% | Transcription, chromatin and RNA processing | | none | 1 of 11 |
| ***Cpeb1*** | A3SS | — | In-frame · Loop/Disordered | 0.103 | +0.146 | 0.007 | -0.197 | 0.003 | -0.051 | 135% | Synaptic signaling / neuronal plasticity | RNA processing and translation | none | 1 of 10 |
| ***Cpeb2*** | SE | 640-669 | In-frame · Loop/Disordered | 0.793 | +0.146 | 0.018 | -0.169 | 0.027 | -0.023 | 116% | Synaptic signaling / neuronal plasticity | RNA processing and translation | none | 1 of 4 |
| ***D16Ertd472e*** | A5SS | — | Poison exon → NMD | 0.623 | +0.321 | <0.001 | -0.206 | 0.002 | +0.115 | 64% | Not assigned | | none | 1 of 3 |
| ***Dctn1*** | SE | 132-138 | In-frame · Loop/Disordered | 0.546 | +0.278 | <0.001 | -0.256 | 0.002 | +0.022 | 92% | Synaptic signaling / neuronal plasticity | Calcium and cytoskeletal signaling | none | 2 of 11 |

| Gene | Event type | Reversed segment (aa) | Structure-function call | Baseline PSI | Cd vs Veh ΔPSI | Cd FDR | Cd+Akk vs Cd ΔPSI | Cd+Akk vs Cd FDR | Cd+Akk vs VehΔPSI | % restoration | Functional group | Figure 7 group | Akk-alone effect | Cd-altered events (of total in gene) |
|---|---|---|---|---|---|---|---|---|---|---|---|---|---|---|
| ***Dtx2*** | RI | — | Poison exon → NMD | 0.438 | +0.423 | <0.001 | -0.307 | 0.002 | +0.115 | 73% | Proteostasis, DNA damage and stress response | Ubiquitin, proteostasis and repair | none | 1 of 2 |
| ***Exosc8*** | SE | 19-39 | Poison exon → NMD | 0.877 | +0.123 | <0.001 | -0.087 | 0.011 | +0.036 | 71% | Transcription, chromatin and RNA processing | | same direction as Cd | 1 of 5 |
| ***Faf2*** | A5SS | — | In-frame · Loop/Disordered | 0.695 | +0.260 | <0.001 | -0.183 | <0.001 | +0.077 | 70% | Proteostasis, DNA damage and stress response | | none | 1 of 5 |
| ***Hdac6*** | A5SS | — | In-frame · Loop/Disordered | 0.201 | +0.466 | <0.001 | -0.302 | 0.047 | +0.165 | 65% | Proteostasis, DNA damage and stress response | Ubiquitin, proteostasis and repair | none | 1 of 5 |
| ***Inpp5k*** | SE | 387-414 | In-frame · Functional-element (GPR78/PAK1 region) | 0.854 | +0.110 | <0.001 | -0.075 | 0.034 | +0.035 | 68% | Metabolism, transport and cell structure | | none | 1 of 16 |
| ***Jak3*** | SE | 564-591 | Poison exon → NMD | 0.356 | +0.166 | 0.048 | -0.218 | 0.010 | -0.052 | 131% | Proteostasis, DNA damage and stress response | | none | 2 of 4 |
| ***Lect1*** | A3SS | — | In-frame · Loop/Disordered | 0.572 | +0.362 | 0.006 | -0.252 | 0.003 | +0.110 | 70% | Metabolism, transport and cell structure | | none | 1 of 1 |
| ***Med12l*** | SE | — | In-frame · Loop/Disordered | 0.548 | +0.452 | <0.001 | -0.370 | 0.009 | +0.082 | 82% | Transcription, chromatin and RNA processing | Transcription and chromatin | none | 1 of 14 |
| ***Mettl10*** | SE | 134-205 | Poison exon → NMD | 0.808 | +0.174 | <0.001 | -0.072 | 0.004 | +0.102 | 41% | Transcription, chromatin and RNA processing | | same direction as Cd | 1 of 9 |
| ***Mthfsl*** | SE | — | Poison exon → NMD | 0.292 | +0.472 | 0.002 | -0.480 | 0.012 | -0.008 | 102% | Metabolism, transport and cell structure | | none | 3 of 11 |
| ***Mynn*** | A5SS | — | Poison exon → NMD | 0.684 | +0.265 | 0.003 | -0.219 | 0.004 | +0.046 | 83% | Transcription, chromatin and RNA processing | | none | 1 of 2 |
| ***Phactr4*** | SE | 64-90 | In-frame · Functional-element | 0.415 | +0.406 | 0.005 | -0.640 | <0.001 | -0.234 | 158% | Synaptic signaling / neuronal plasticity | Calcium and cytoskeletal signaling | none | 1 of 1 |
| ***Phrf1*** | SE | 487-538 | Poison exon → NMD | 0.854 | +0.112 | 0.002 | -0.078 | 0.009 | +0.035 | 70% | Transcription, chromatin and RNA processing | | same direction as Cd | 1 of 6 |
| ***Pias3*** | A5SS | — | Poison exon → NMD | 0.890 | +0.102 | <0.001 | -0.065 | 0.009 | +0.037 | 64% | Proteostasis, DNA damage and stress response | | same direction as Cd | 1 of 3 |
| ***Prdm5*** | SE | 258-284 | Poison exon → NMD | 0.594 | +0.322 | 0.007 | -0.346 | 0.003 | -0.025 | 107% | Transcription, chromatin and RNA processing | Transcription and chromatin | none | 1 of 4 |
| ***Proca1*** | SE | 55-98 | Poison exon → NMD | 0.092 | +0.666 | 0.003 | -0.595 | 0.010 | +0.072 | 89% | Not assigned | | none | 1 of 2 |

| Gene | Event type | Reversed segment (aa) | Structure-function call | Baseline PSI | Cd vs Veh ΔPSI | Cd FDR | Cd+Akk vs Cd ΔPSI | Cd+Akk vs Cd FDR | Cd+Akk vs VehΔPSI | % restoration | Functional group | Figure 7 group | Akk-alone effect | Cd-altered events (of total in gene) |
|---|---|---|---|---|---|---|---|---|---|---|---|---|---|---|
| ***Ptpn2*** | SE | 54-87 | Poison exon → NMD | 0.729 | +0.213 | <0.001 | -0.129 | 0.005 | +0.084 | 61% | Proteostasis, DNA damage and stress response | | same direction as Cd | 1 of 5 |
| ***Rad52*** | SE | 157-182 | Poison exon → NMD | 0.537 | +0.286 | <0.001 | -0.224 | <0.001 | +0.062 | 78% | Proteostasis, DNA damage and stress response | | none | 3 of 11 |
| ***Rnf32*** | SE | 94-141 | Poison exon → NMD | 0.831 | +0.124 | 0.006 | -0.070 | 0.042 | +0.054 | 56% | Proteostasis, DNA damage and stress response | | none | 1 of 7 |
| ***Ryr3*** | A3SS | — | In-frame · Loop/Disordered | 0.828 | +0.146 | 0.016 | -0.211 | 0.008 | -0.064 | 145% | Synaptic signaling / neuronal plasticity | Calcium and cytoskeletal signaling | none | 1 of 19 |
| ***Sag*** | SE | 338-345 | In-frame · Functional-element | 0.702 | +0.269 | 0.012 | -0.222 | 0.004 | +0.047 | 83% | Synaptic signaling / neuronal plasticity | | none | 1 of 2 |
| ***Syce2*** | A3SS | — | In-frame · Loop/Disordered | 0.000 | +0.197 | 0.003 | -0.108 | 0.008 | +0.089 | 55% | Not assigned | | none | 1 of 2 |
| ***Tanc2*** | SE | — | In-frame · Loop/Disordered | 0.083 | +0.267 | 0.006 | -0.240 | 0.020 | +0.027 | 90% | Synaptic signaling / neuronal plasticity | | none | 1 of 6 |
| ***Tie1*** | RI | — | Poison exon → NMD | 0.088 | +0.221 | 0.010 | -0.210 | 0.016 | +0.012 | 95% | Metabolism, transport and cell structure | | none | 1 of 4 |
| ***Trp63*** | A5SS | — | In-frame · Loop/Disordered | 0.191 | +0.679 | <0.001 | -0.512 | 0.028 | +0.167 | 75% | Transcription, chromatin and RNA processing | Transcription and chromatin | none | 1 of 2 |
| ***Usp40*** | MXE | — | Poison exon → NMD | 0.173 | +0.451 | 0.002 | -0.603 | <0.001 | -0.152 | 134% | Proteostasis, DNA damage and stress response | Ubiquitin, proteostasis and repair | none | 2 of 7 |
| ***Zfand2b*** | RI | — | Poison exon → NMD | 0.165 | +0.109 | 0.008 | -0.142 | <0.001 | -0.033 | 130% | Proteostasis, DNA damage and stress response | | none | 1 of 3 |
| ***Zfp518b*** | SE | — | In-frame · Loop/Disordered | 0.064 | +0.500 | 0.003 | -0.408 | 0.043 | +0.092 | 82% | Transcription, chromatin and RNA processing | Transcription and chromatin | none | 1 of 2 |
| ***Cd decreased inclusion (n = 50)*** | | | | | | | | | | | | | | |
| ***1700001L05Rik*** | SE | — | Poison exon → NMD | 0.875 | -0.293 | 0.023 | +0.418 | <0.001 | +0.125 | 143% | Not assigned | | none | 1 of 2 |
| ***2210408F21Rik*** | SE | — | Poison exon → NMD | 1.000 | -0.437 | <0.001 | +0.349 | <0.001 | -0.088 | 80% | Not assigned | | none | 6 of 19 |
| ***5730480H06Rik*** | SE | — | In-frame · Loop/Disordered | 0.988 | -0.154 | 0.001 | +0.156 | 0.002 | +0.002 | 101% | Not assigned | | none | 1 of 4 |
| ***A230009B12Rik*** | SE | — | In-frame · Loop/Disordered | 0.879 | -0.590 | <0.001 | +0.462 | 0.001 | -0.128 | 78% | Not assigned | | none | 1 of 1 |
| ***Arntl2*** | A3SS | — | Poison exon → NMD | 0.403 | -0.346 | <0.001 | +0.179 | 0.024 | -0.167 | 52% | Transcription, chromatin and RNA processing | | same direction as Cd | 1 of 2 |

| Gene | Event type | Reversed segment (aa) | Structure-function call | Baseline PSI | Cd vs Veh ΔPSI | Cd FDR | Cd+Akk vs Cd ΔPSI | Cd+Akk vs Cd FDR | Cd+Akk vs VehΔPSI | % restoration | Functional group | Figure 7 group | Akk-alone effect | Cd-altered events (of total in gene) |
|---|---|---|---|---|---|---|---|---|---|---|---|---|---|---|
| ***Bap1*** | RI | — | Poison exon → NMD | 0.947 | -0.131 | 0.016 | +0.162 | 0.017 | +0.031 | 124% | Proteostasis, DNA damage and stress response | | none | 1 of 17 |
| ***Bco2*** | SE | 126-164 | Poison exon → NMD | 0.962 | -0.315 | 0.021 | +0.353 | 0.019 | +0.038 | 112% | Metabolism, transport and cell structure | | none | 1 of 2 |
| ***Cbx7*** | A3SS | — | In-frame · Loop/Disordered | 0.952 | -0.114 | 0.010 | +0.121 | 0.031 | +0.007 | 106% | Transcription, chromatin and RNA processing | | same direction as Cd | 1 of 3 |
| ***Ccdc13*** | SE | 500-527 | In-frame · Functional-element (SCF/Cdc4 degron) | 1.000 | -0.136 | 0.004 | +0.116 | 0.011 | -0.020 | 85% | Metabolism, transport and cell structure | | none | 1 of 2 |
| ***Ccdc77*** | A3SS | — | In-frame · Loop/Disordered | 0.425 | -0.393 | 0.003 | +0.520 | 0.004 | +0.127 | 132% | Metabolism, transport and cell structure | Cilia and centrosome | none | 1 of 9 |
| ***Cemip*** | SE | 1136-1204 | Poison exon → NMD | 0.985 | -0.110 | 0.025 | +0.103 | 0.042 | -0.007 | 94% | Metabolism, transport and cell structure | | none | 1 of 2 |
| ***Cfap69*** | SE | 229-287 | Poison exon → NMD | 1.000 | -0.155 | 0.008 | +0.122 | 0.040 | -0.033 | 79% | Metabolism, transport and cell structure | Cilia and centrosome | none | 2 of 21 |
| ***Crem*** | MXE | — | In-frame · Loop/Disordered | 0.495 | -0.327 | <0.001 | +0.313 | <0.001 | -0.013 | 96% | Transcription, chromatin and RNA processing | Transcription and chromatin | none | 1 of 20 |
| ***Csgalnact1*** | SE | — | Poison exon → NMD | 1.000 | -0.205 | 0.002 | +0.121 | 0.017 | -0.084 | 59% | Metabolism, transport and cell structure | | none | 1 of 5 |
| ***Ctbs*** | SE | — | Poison exon → NMD | 1.000 | -0.322 | 0.001 | +0.218 | 0.020 | -0.103 | 68% | Metabolism, transport and cell structure | | none | 1 of 2 |
| ***Ctns*** | SE | 228-284 | In-frame · Loop/Disordered | 0.056 | -0.101 | 0.003 | +0.124 | 0.001 | +0.023 | 123% | Metabolism, transport and cell structure | Calcium and ion transport | none | 2 of 6 |
| ***Cyb561d1*** | SE | 50-62 | Poison exon → NMD | 1.000 | -0.191 | 0.015 | +0.167 | 0.036 | -0.024 | 87% | Metabolism, transport and cell structure | | none | 1 of 4 |
| ***Efna5*** | SE | 162-188 | In-frame · Loop/Disordered | 0.931 | -0.284 | 0.018 | +0.342 | <0.001 | +0.058 | 120% | Synaptic signaling / neuronal plasticity | Axon guidance and synapse formation | none | 1 of 1 |
| ***Eif5a*** | SE | — | Poison exon → NMD | 0.971 | -0.126 | <0.001 | +0.126 | <0.001 | +0.000 | 100% | Transcription, chromatin and RNA processing | | none | 1 of 9 |
| ***Erc1*** | SE | — | Poison exon → NMD | 0.934 | -0.106 | 0.024 | +0.115 | 0.028 | +0.009 | 108% | Synaptic signaling / neuronal plasticity | | none | 1 of 11 |
| ***Exosc5*** | A3SS | — | Poison exon → NMD | 0.196 | -0.135 | <0.001 | +0.095 | 0.014 | -0.041 | 70% | Transcription, chromatin and RNA processing | | none | 1 of 2 |
| ***Fbf1*** | SE | — | In-frame · Loop/Disordered | 0.646 | -0.536 | 0.003 | +0.439 | 0.041 | -0.098 | 82% | Metabolism, transport and cell structure | Cilia and centrosome | none | 1 of 19 |
| ***Fndc1*** | SE | 391-453 | In-frame · Removes domain (Fibronectin III domain) | 0.898 | -0.234 | 0.018 | +0.163 | 0.041 | -0.071 | 70% | Metabolism, transport and cell structure | | none | 1 of 2 |

| Gene | Event type | Reversed segment (aa) | Structure-function call | Baseline PSI | Cd vs Veh ΔPSI | Cd FDR | Cd+Akk vs Cd ΔPSI | Cd+Akk vs Cd FDR | Cd+Akk vs VehΔPSI | % restoration | Functional group | Figure 7 group | Akk-alone effect | Cd-altered events (of total in gene) |
|---|---|---|---|---|---|---|---|---|---|---|---|---|---|---|
| ***Garnl3*** | A5SS | — | Poison exon → NMD | 0.978 | -0.106 | 0.002 | +0.083 | 0.014 | -0.024 | 78% | Synaptic signaling / neuronal plasticity | | none | 2 of 12 |
| ***Gm20498*** | A5SS | — | In-frame · Removes domain | 0.806 | -0.532 | 0.004 | +0.631 | <0.001 | +0.099 | 119% | Not assigned | | none | 1 of 6 |
| ***Gm2762*** | SE | — | In-frame · Loop/Disordered | 1.000 | -0.533 | <0.001 | +0.374 | 0.004 | -0.159 | 70% | Not assigned | | none | 1 of 10 |
| ***Gtpbp6*** | A3SS | — | Poison exon → NMD | 0.941 | -0.158 | 0.039 | +0.153 | 0.017 | -0.005 | 97% | Metabolism, transport and cell structure | | none | 1 of 4 |
| ***Hdac4*** | SE | — | Poison exon → NMD | 0.945 | -0.118 | 0.007 | +0.141 | <0.001 | +0.024 | 119% | Transcription, chromatin and RNA processing | Transcription and chromatin | none | 1 of 5 |
| ***Intu*** | SE | 232-260 | Poison exon → NMD | 0.930 | -0.283 | 0.002 | +0.353 | <0.001 | +0.070 | 125% | Metabolism, transport and cell structure | Cilia and centrosome | none | 1 of 2 |
| ***Lrrc9*** | SE | 1368-1391 | In-frame · Loop/Disordered | 0.956 | -0.592 | <0.001 | +0.279 | 0.004 | -0.313 | 47% | Not assigned | | none | 1 of 2 |
| ***Lyrm1*** | SE | — | In-frame · Loop/Disordered | 0.932 | -0.382 | 0.026 | +0.324 | 0.021 | -0.058 | 85% | Metabolism, transport and cell structure | | none | 1 of 7 |
| ***Magt1*** | RI | — | Poison exon → NMD | 0.936 | -0.254 | <0.001 | +0.282 | <0.001 | +0.028 | 111% | Metabolism, transport and cell structure | Calcium and ion transport | none | 1 of 1 |
| ***Mfsd11*** | A3SS | — | Poison exon → NMD | 0.986 | -0.150 | 0.010 | +0.135 | 0.011 | -0.014 | 90% | Metabolism, transport and cell structure | Calcium and ion transport | none | 1 of 7 |
| ***Ncoa6*** | SE | 1970-1992 | In-frame · Functional-element (EP300 region) | 0.515 | -0.376 | 0.021 | +0.327 | 0.041 | -0.049 | 87% | Transcription, chromatin and RNA processing | Transcription and chromatin | none | 1 of 4 |
| ***Ogg1*** | A3SS | — | Poison exon → NMD | 0.948 | -0.101 | 0.027 | +0.153 | 0.014 | +0.052 | 151% | Proteostasis, DNA damage and stress response | | none | 1 of 5 |
| ***Pak3*** | SE | — | In-frame · Loop/Disordered | 0.311 | -0.242 | <0.001 | +0.099 | 0.005 | -0.143 | 41% | Synaptic signaling / neuronal plasticity | Axon guidance and synapse formation | none | 1 of 7 |
| ***Plpp5*** | SE | — | Poison exon → NMD | 0.142 | -0.110 | <0.001 | +0.047 | 0.034 | -0.063 | 43% | Metabolism, transport and cell structure | | none | 1 of 6 |
| ***Plxnb3*** | A3SS | 5-314 | In-frame · Removes domain | 0.977 | -0.172 | 0.005 | +0.178 | 0.005 | +0.006 | 103% | Synaptic signaling / neuronal plasticity | Axon guidance and synapse formation | none | 1 of 10 |
| ***Ppcdc*** | SE | 121-176 | Poison exon → NMD | 0.954 | -0.243 | <0.001 | +0.289 | <0.001 | +0.046 | 119% | Metabolism, transport and cell structure | | none | 1 of 2 |
| ***Ralgps1*** | A3SS | — | In-frame · Loop/Disordered | 0.951 | -0.269 | 0.002 | +0.141 | 0.034 | -0.128 | 52% | Metabolism, transport and cell structure | | none | 1 of 14 |
| ***Ryr1*** | SE | 3868-3873 | In-frame · Loop/Disordered | 0.976 | -0.133 | 0.038 | +0.157 | 0.020 | +0.024 | 118% | Synaptic signaling / neuronal plasticity | Calcium and ion transport | none | 1 of 5 |
| ***Saal1*** | A3SS | — | Poison exon → NMD | 0.956 | -0.126 | 0.012 | +0.130 | 0.001 | +0.004 | 103% | Metabolism, transport and cell structure | | none | 1 of 4 |
| ***Sema4a*** | A3SS | — | Poison exon → NMD | 0.148 | -0.108 | 0.041 | +0.109 | 0.038 | +0.001 | 101% | Synaptic signaling / neuronal plasticity | Axon guidance and synapse formation | none | 2 of 10 |

| Gene | Event type | Reversed segment (aa) | Structure-function call | Baseline PSI | Cd vs Veh ΔPSI | Cd FDR | Cd+Akk vs Cd ΔPSI | Cd+Akk vs Cd FDR | Cd+Akk vs VehΔPSI | % restoration | Functional group | Figure 7 group | Akk-alone effect | Cd-altered events (of total in gene) |
|---|---|---|---|---|---|---|---|---|---|---|---|---|---|---|
| ***Sgk3*** | RI | — | Poison exon → NMD | 0.111 | -0.102 | <0.001 | +0.144 | <0.001 | +0.043 | 141% | Metabolism, transport and cell structure | | none | 1 of 6 |
| ***Smc6*** | SE | 86-120 | Poison exon → NMD | 1.000 | -0.197 | <0.001 | +0.150 | 0.005 | -0.046 | 76% | Proteostasis, DNA damage and stress response | | same direction as Cd | 3 of 7 |
| ***Smim1*** | A3SS | — | Poison exon → NMD | 0.746 | -0.487 | 0.006 | +0.442 | 0.003 | -0.045 | 91% | Not assigned | | none | 2 of 9 |
| ***Tcf7l2*** | A5SS | — | In-frame · Loop/Disordered | 0.965 | -0.380 | 0.047 | +0.371 | 0.008 | -0.009 | 98% | Transcription, chromatin and RNA processing | Transcription and chromatin | none | 1 of 11 |
| ***Trmt13*** | A3SS | — | Poison exon → NMD | 0.655 | -0.175 | 0.001 | +0.219 | <0.001 | +0.044 | 125% | Transcription, chromatin and RNA processing | | none | 2 of 6 |
| ***Upp2*** | SE | 132-169 | Poison exon → NMD | 0.950 | -0.275 | 0.004 | +0.259 | 0.005 | -0.016 | 94% | Metabolism, transport and cell structure | | none | 1 of 3 |
| ***Zbtb17*** | RI | — | In-frame · Removes domain | 1.000 | -0.117 | 0.002 | +0.098 | 0.007 | -0.019 | 84% | Transcription, chromatin and RNA processing | | none | 1 of 6 |

| Supplementary Table S2. mRNA expression-anchored data integration with $m^6A$ and AS | | | |
|---|---|---|---|
| **Analysis** | **Cd vs Vehicle** | **Akk vs Vehicle** | **Cd+Akk vs Cd** |
| **$m^6A$ epitranscriptomics** | | | |
| Significant $m^6A$ events, $p < 0.05$ | 402 | 1,075 | 1,536 |
| Unique genes with significant $m^6A$ changes | 377 | 979 | 1,425 |
| Genes also significant by RNA-seq, $p < 0.05$ | 56 | 74 | 26 |
| % of $m^6A$ genes also showing mRNA abundance change | 14.9% | 7.6% | 1.8% |
| Enrichment of RNA-seq overlap over random expectation | 3.3-fold | 2.1-fold | 1.2-fold |
| **Alternative splicing (AS)** | | | |
| Significant AS events, $p < 0.05$ | 4,128 | 2,960 | 3,571 |
| Significant AS events, FDR < 0.05 | 1,104 | 672 | 754 |
| Unique genes with significant AS | 2,802 | 2,140 | 2,486 |
| Unique genes with significant AS, FDR < 0.05 | 938 | 573 | 644 |
| Genes also significant by RNA-seq, $p < 0.05$ | 167 | 107 | 33 |
| % of AS genes also showing mRNA abundance change | 6.0% | 5.0% | 1.3% |
| Enrichment of RNA-seq overlap over random expectation | 1.0-fold | 0.9-fold | 0.7-fold |

**Supplementary Table S3. Gene-set membership and per-layer significance thresholds (Figure 8).**

**A. Gene sets evaluated in Figure 8.**

| Gene set | Genes (n) | Figure 8 panel | Source |
|---|---|---|---|
| $m^6A$ regulators | 22 | A (left ring of each pair) | Curated mouse $m^6A$ writers, readers and erasers |
| Alternative-splicing regulators | 72 | A (right ring of each pair) | Curated mouse splicing regulatory RNA-binding proteins, comprising the SR and hnRNP families and other characterized splicing factors |
| Learning and memory | 72 | contributes to B | GO Biological Process 2025: learning, memory and long-term memory |
| Alzheimer disease | 161 | contributes to B | KEGG 2019 Mouse: Alzheimer disease |
| Combined cognitive and Alzheimer-disease set, nonduplicated | 227 | B | Union of the two sets above. Six genes belong to both (Apoe, App, Mapt, Plcb1, Ppp3cb, Psen1) and are counted once |

**B. Per-layer significance thresholds.**

| Regulatory layer | Data and method | Threshold |
|---|---|---|
| Alternative splicing | rMATS (JCEC); SE, A5SS, A3SS, MXE and RI events | $p < 0.05$ |
| $m^6A$ modification | Arraystar Mouse Epitranscriptomic Microarray | $p < 0.05$ |
| mRNA abundance | RNA-seq (Cuffdiff) | $p < 0.05$ |

**C. Gene-set membership.**

**$m^6A$ regulators (n = 22).** Mettl3, Mettl14, Wtap, Virma, Rbm15, Rbm15b, Zc3h13, Mettl16, Ythdf1, Ythdf2, Ythdf3, Ythdc1, Ythdc2, Igf2bp1, Igf2bp2, Igf2bp3, Hnrnpa2b1, Hnrnpc, Rbmx, Prrc2a, Fto, Alkbh5

**Alternative-splicing regulators (n = 72).** Srsf1, Srsf2, Srsf3, Srsf4, Srsf5, Srsf6, Srsf7, Srsf9, Srsf10, Srsf11, Srsf12, Srrm1, Srrm2, Srrm4, Tra2a, Tra2b, Hnrnpa1, Hnrnpa2b1, Hnrnpa3, Hnrnpb1, Hnrnpc, Hnrnpd, Hnrnpdl, Hnrnpf, Hnrnph1, Hnrnph2, Hnrnph3, Hnrnpk, Hnrnpl, Hnrnpll, Hnrnpm, Hnrnpu, Hnrnpul1, Ptbp1, Ptbp2, Ptbp3, Rbfox1, Rbfox2, Rbfox3, Mbnl1, Mbnl2, Mbnl3, Celf1, Celf2, Celf3, Celf4, Celf5, Celf6, Esrp1, Esrp2, Nova1, Nova2, Khdrbs1, Khdrbs2, Khdrbs3, Qki, Tia1, Tial1, Rbm4, Rbm10, Rbm20, Rbm24, Rbm38, Rbm39, Rbm47, Sf1, U2af1, U2af2, Khsrp, Fmr1, Msi1, Msi2

**Combined cognitive and Alzheimer-disease set (n = 227).** Acss2, Adam10, Adam17, Adcy8, Apaf1, Apbb1, Aph1a, Aph1b, Aph1c, Apoe, App, Arc, Atad1, Atf6, Atp2a1, Atp2a2, Atp2a3, Atp5a1, Atp5b, Atp5c1, Atp5d, Atp5e, Atp5f1, Atp5g1, Atp5g2, Atp5g3, Atp5h, Atp5j, Atp5o, Atxn1, Atxn1l, Bace1, Bace2, Bad, Bid, Brsk1, Cacna1c, Cacna1d, Cacna1f, Cacna1s, Calb1, Calm1, Calm2, Calm3, Calml4, Camk2n1, Camk4, Capn1, Capn2, Casp12, Casp3, Casp7, Casp8, Casp9, Cdk5, Cdk5r1, Chrnb2, Cic, Cldn5, Cln3, Cntnap2, Cox4i1, Cox4i2, Cox5a, Cox5b, Cox6a1, Cox6a2, Cox6b1, Cox6b2, Cox6c, Cox7a1, Cox7a2, Cox7a2l, Cox7b, Cox7b2, Cox7c, Cox8a, Cox8b, Cpeb3, Cux2, Cyc1, Cycs, Drd2, Drd3, Eif2ak3, Eif2ak4, Eif4ebp2, Ephb2, Ern1, Fadd, Fas, Fgf13, Fyn, Gapdh, Gapdh-ps15, Gnaq, Gpr88, Grin1, Grin2a, Grin2b, Grin2c, Grin2d, Gsk3b, Hsd17b10, Ide, Il1b, Insr, Itpr1, Itpr2, Itpr3, Kat2a, Klk8, Ldlr, Lgmn, Llgl1, Lpl, Lrp1, Lrrn4, Mapk1, Mapk3, Mapk8ip2, Mapt, Metap1, Mme, Musk, Nae1, Ncstn, Ndufa1, Ndufa10, Ndufa11, Ndufa12, Ndufa13, Ndufa2, Ndufa3, Ndufa4, Ndufa4l2, Ndufa5, Ndufa6, Ndufa7, Ndufa8, Ndufa9, Ndufab1, Ndufb10, Ndufb11, Ndufb2, Ndufb3, Ndufb4, Ndufb5,

Ndufb6, Ndufb7, Ndufb8, Ndufb9, Ndufc1, Ndufc2, Ndufs1, Ndufs2, Ndufs3, Ndufs4, Ndufs5, Ndufs6, Ndufs7, Ndufs8, Ndufv1, Ndufv2, Ndufv3, Neto1, Nfatc4, Nlgn3, Nos1, Npas4, Nrxn1, Nrxn3, Ntrk2, Pja2, Plcb1, Plcb2, Plcb3, Plcb4, Plk2, Ppp3ca, Ppp3cb, Ppp3cc, Ppp3r1, Ppp3r2, Prnp, Psen1, Psen2, Psenen, Ptn, Rasgrf1, Reln, Rgs14, Rtn3, Rtn4, Ryr3, Sdha, Sdhb, Sdhc, Sdhd, Sgk1, Shank1, Shank2, Shank3, Shisa7, Slc2a4, Slc6a1, Slc8a2, Slc8a3, Snca, Stra6, Synj1, Syt12, Th, Tlr2, Tnf, Tnfrsf1a, Trem2, Uqcr10, Uqcr11, Uqcrb, Uqcrc1, Uqcrc2, Uqcrfs1, Uqcrh, Uqcrq, Vldlr, Ythdf1

*A nominal threshold was applied consistently across all three layers, as described in Materials and Methods: multiple-testing correction is highly stringent in high-dimensional m⁶A and splicing data, where thousands of transcript- or event-level features are tested, and would exclude moderate changes that become informative when read across layers. Where Benjamini–Hochberg adjustment was applied instead — the differentially expressed gene sets in Figure 2A, the enrichment analyses in Figure 3, the 12-transcript reversal set in Figure 4A, and the high-confidence splicing set in Figure 7 and Supplementary Table S1 — it is stated in the corresponding legend. The number of affected genes behind every ring is printed inside that ring in Figure 8. Learning and memory and Alzheimer-disease symbols are returned by GO and KEGG in human uppercase and are listed above in mouse nomenclature.*
